# Comparative study of protein X-ray and NMR structures: molecular docking-based virtual screening

**DOI:** 10.1101/2025.03.25.645374

**Authors:** Srdan Masirevic, Hao Fan

## Abstract

Molecular docking-based virtual ligand screening is a powerful computational approach for identifying potential binders from large chemical libraries. Protein structures used in docking screens are commonly derived from X-ray crystallography or NMR spectroscopy, yet their impact on screening performance remains unclear. To address this, we conducted virtual screening using Glide against both apo and holo X-ray and NMR structures of 18 proteins. While no statistically significant difference in screening performance was observed for apo structures overall, X-ray apo structures tended to perform better in cases where better ligand enrichment than random selection was achieved. Similarly, single holo X-ray and NMR structures did not exhibited statistically significant difference in screening performance either. However, when multiple holo X-ray structures and NMR conformers (from one PDB ensemble) per protein were used, X-ray structures outperformed NMR conformers in most cases. In addition, for consensus enrichment which leverages multiple structures/conformers per protein to optimise ligand ranking, X-ray holo structures exhibited better performance than NMR holo conformers, suggesting that X-ray structures with chemically diverse co-crystalized ligands may introduce more relevant binding-site configurations than the NMR conformers with higher structural diversity but the same bound ligand. Overall, the better performance by X-ray holo structures could be partially attributed to the higher numbers of hydrogen bonds and hydrophobic contacts, formed between proteins and docked ligands.

## 1. Introduction

Employment of computational techniques to study protein-ligand interactions has significantly increased in recent years, as they help to accelerate the discovery of small molecules for various purposes (1, 2). In particular, Molecular docking-based virtual ligand screening has become a standard method used to predict new ligands from large libraries of small molecules (3–7). Each library molecule is docked into a binding site of the target protein structure, then scored and ranked based on its complementarity to the protein. Typically, top-ranked docked molecules are selected for experimental validation. Performance of docking screens represented by prioritising binders over non-binders is often evaluated using metrics like the area under the curve of enrichment plot (logAUC) with x-axis in logarithmic scale, which emphasises the early ligand enrichment (8, 9).

X-ray crystallography and NMR spectroscopy are commonly used experimental techniques for determining structural and functional features of proteins at the atomic level. X-ray crystallography can capture high resolution 3D snapshots of proteins in the crystal lattice and can be applied to proteins with high molecular weight (>500 kDa) (10). On the other hand, NMR spectroscopy, excels at revealing the dynamic properties of proteins and their physiologically relevant conformations in solution (11). To date, both types of structures have successfully contributed to the discovery of new ligands (12, 13). However, given the differences between both techniques, it is generally accepted that the structure determined in solution will not be identical to the one influenced by the packing of the atoms in the crystal lattice. Hence, previous studies have examined structural differences between X-ray and NMR protein structures on a large scale. Garbuzinskiy et al. (14) found that NMR structures have more contacts per residue at the distances < 3 Å, and between 4.5-6.5 Å, but fewer contacts at distances between 3.0-4.5 Å and 6.5-8.0 Å, while Melnik et al. (15) noted weak similarity in hydrogen bonding patterns (R = 0.32) between the two experimental structures, with X-ray structures generally containing higher number of hydrogen bonds in the main chain. Sikic et al. (16) conducted a systematic comparison of X-ray and NMR structures highlighting that amino acid side-chain pairs, particularly those that are hydrophilic, often adopt different orientations. Schneider et al. (17) proposed that X-ray structures generally offer better templates for protein design due to the reduced variability in side-chain conformations, while NMR conformers shall be used in the absence of high resolution X-ray structure.

Given these structural differences between X-ray and NMR structures, it is expected that virtual screening performance could be affected by whether X-ray or NMR structures are used, as even slight discrepancies in atomic positions can affect virtual screening performance. For instance, the difference in hydrogen bonding patterns and contacts per residue observed in X-ray versus NMR structures (14, 15) may dictate the availability of residues for ligand interactions, ultimately influencing virtual screening performance.

In this study, we aimed to address the following questions: 1. Which type of protein structure — X-ray or NMR in the ligand-free (apo) or ligand-bound (holo) state — is more suitable for ligand identification using molecular docking-based virtual screening? 2. Does docking screens against multiple X-ray structures or NMR conformers of the same protein deliver improved performance compared to using a single structure? 3. If one type of structure yields better virtual screening performance, can this be explained or at least correlated to specific protein/ligand features?

To address these questions, we assembled a library of 18 proteins, each of which has both X-ray and NMR structures, and conducted molecular docking-based virtual screening utilising the Glide software. The virtual screening performance was evaluated using ligand enrichment (logAUC). For apo structures, in cases where ligand enrichment was above the random selection threshold (logAUC > 14.5) for both structures, X-ray apo structures tended to perform better than NMR apo structures, though the difference remained statistically insignificant. For single holo structures, virtual screening performance was not statistically different between X-ray and NMR structures either. However, when multiple holo structures/ conformers of the target protein were considered, X-ray structures from different PDB entries generally outperformed NMR conformers from the same PDB entry, with statistical significance observed across all pairwise comparisons.

To understand the factors influencing these differences, we analysed protein and ligand features. We found that differences in binding pocket features, such as volume, solvent-accessible surface area (SASA), hydrophobicity, and the proportion of polar atoms, between X-ray and NMR structures showed weak correlations with differences in virtual screening performance between the two structure types (R² ≤ 0.21). However, differences in protein-ligand hydrogen bonds and hydrophobic contacts between the two structure types showed higher correlations (R² = 0.43 and R² = 0.35, respectively) with ligand enrichment differences. Additionally, NMR holo conformers from one PDB entry exhibited greater structural deviations in the binding pocket (heavy atoms RMSD ranging from 1.46 Å to 2.02 Å) compared to that of X-ray holo structures from different PDB entries ( heavy atoms RMSD ranging from 0.42 Å to 1.77 Å), suggesting greater structural heterogeneity in the binding pocket of conformers in the same NMR ensemble. These findings suggest that incorporating experimental structures that capture diverse ligand-binding motifs may be more beneficial for docking screens, than simply increasing binding site conformational diversity.

## 2. Results and Discussions

### 2.1 Cognate docking and geometry assessment

To evaluate the robustness of the docking protocol in reproducing experimental ligand binding motifs, we redocked each cognate ligand into its native holo X-ray and holo NMR structure and calculated the RMSD between the docked pose and the experimentally determined ligand structure. Docking poses with RMSD values of less than 3Å from the experimental ligand structure were considered successful (near-native). Since NMR structures are composed of ensembles of conformers solved in complexes with the same ligand, we compared the performance from the X-ray structure of the best resolution and that from the first conformer in the NMR ensemble. The results of these calculations are presented in Table S1.

In 9 out of 11 (82%) cases of X-ray structures and in 8 out of 11 (73%) cases of NMR conformers, the best-scored docking pose accurately reproduced the experimental ligand geometry. The complexity of the cognate ligand strongly affects the accuracy of ligand docking pose (Figure 1). For instance, the crystal ligand structure with 9 rotatable bonds in Bcl-2 was accurately reproduced (Figure 1A), while the NMR ligand structure with 13 rotatable bonds was not (Figure 1B).

**Figure 1.**
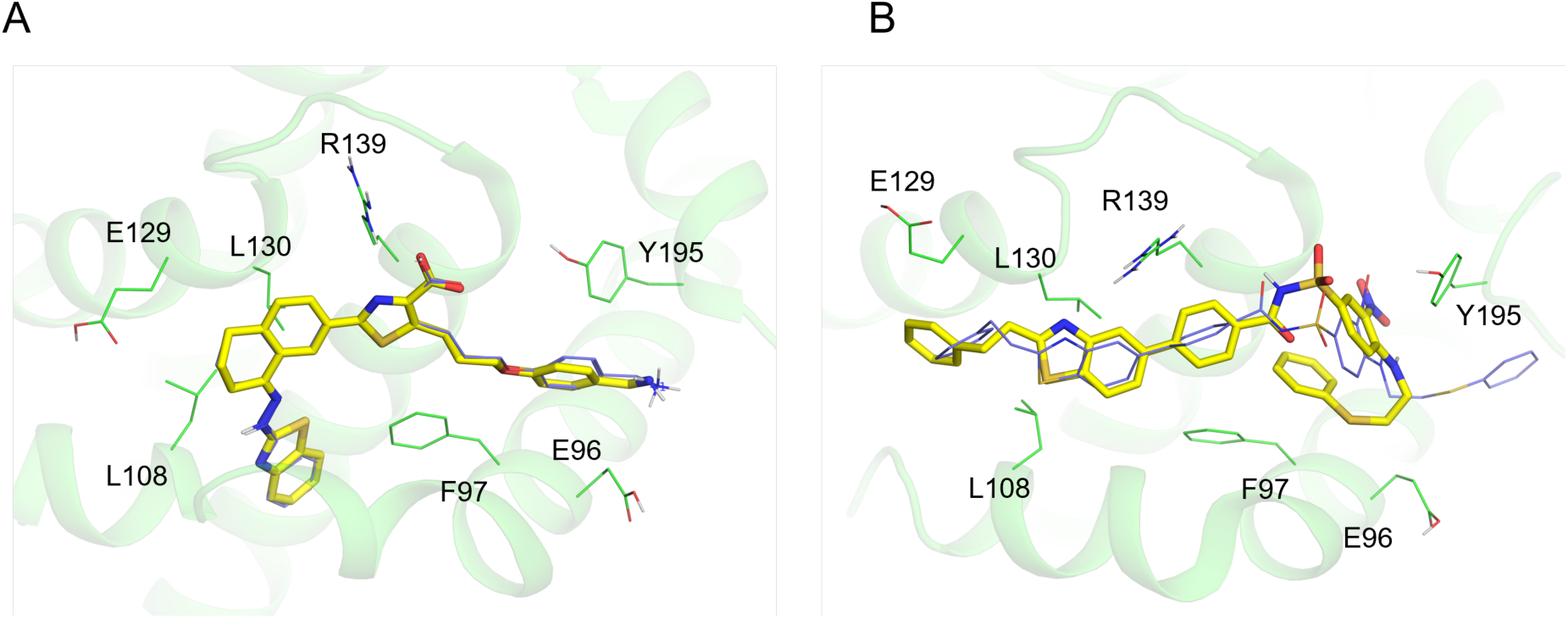
Representative example of the cognate ligand geometry assessment for the Bcl-2 holo structures. **A.** Geometry assessment of Bcl-2 X-ray ligand structure (PDB ID: 3ZLR) (RMSD = 0.30Å). **B**. Geometry assessment of Bcl-2 NMR ligand structure (PDB ID: 2O1Y) (RMSD = 4.15Å). The experimental ligand structure is presented with purple lines, while the ligand docking pose is presented with yellow sticks.

### 2.2 Ligand enrichment of apo structures

To evaluate virtual screening performances of both structure types, we docked each molecule from the virtual screening library to X-ray and NMR apo structures of 12 proteins, and calculated logAUC values of the ligand enrichment plots (Table 2, Figure S1). Among these 12 proteins, ligand enrichment (logAUC) of X-ray apo structure was better, comparable, and worse than that of NMR apo structure in 42%, 25%, and 33% of cases, respectively. However, three proteins: HSP90, Integrin alpha-L and Stromelysin-1 failed to effectively distinguish true binders/ligands from non-binders/decoys, as reflected in their poor ligand enrichments from both X-ray and NMR structures. When these three proteins are excluded, X-ray apo structures performed better, comparable, and worse than NMR apo structures in 56%, 33%, and 11% of cases, respectively, indicating that X-ray apo structures may generally outperform NMR apo structures in virtual screening. Besides this arbitrary comparison, however, statistical analysis of logAUC values across 12 proteins indicates no significant difference between the two structure types (p = 0.460).

**Table 1:** Selected protein targets used in this study.

|  | Protein name<br>(*Species*) | Uniprot<br>ID | PDB ID<br>X-ray (ligand<br>ID) | Resoluti<br>on (Å) | PDB ID NMR<br>(ligand ID) | Number<br>of NMR<br>conform<br>ers |
| --- | --- | --- | --- | --- | --- | --- |
| 7 proteins with only apo structures |  |  |  |  |  |  |
| 1 | Cellular tumor<br>antigen p53<br>(*Homo sapiens*) | P04637 | 2J1X | 1.7 | 2FEJ | 36 |
| 2 | Guanyl-specific<br>ribonuclease T1<br>(*Aspergillus oryzae*) | P00651 | 3RNT | 1.8 | 1YGW | 34 |
| 3 | Replication protein A<br>(70 kDa DNA-binding subunit, REPA)<br>(*Homo sapiens*) | P27694 | 4IPC | 1.2 | 1EWI | 1 |
| 4 | Growth factor receptor-bound protein 2 (GRB2)<br>(*Homo sapiens*) | P62993 | 1GRI | 2.8 | 1FHS | 18 |
| 5 | Heat shock protein (HSP 90-alpha)<br>(*Homo sapiens*) | P07900 | 3T0H | 1.2 | 2K5B | 10 |
| 6 | Integrin alpha-L<br>(*Homo sapiens*) | P20701 | 1LFA | 1.8 | 1DGQ | 22 |
| 7 | Fatty acid-binding protein (FABP)<br>(*Homo sapiens*) | P05413 | 3RSW | 2.6 | 1G5W | 20 |
| 5 proteins have both apo and holo structures |  |  |  |  |  |  |
| 8 | Bcl-2-like protein 1<br>(*Homo sapiens*) | Q07817 | 2P1L<br>3ZLR (X0B) | 2.5<br>2.0 | 2LPC<br>2O1Y (43B) | 21<br>1 |
| 9 | Stromelysin-1<br>(*Homo sapiens*) | P08254 | 1CQR<br>1CIZ (DPS) | 2.0<br>1.6 | 1OO9<br>2JT6 (JT6) | 1<br>1 |
| 10 | Macrophage metalloelastase<br>(*Homo sapiens*) | P39900 | 2OXU<br>3F17 (HS4) | 1.2<br>1.2 | 2KRJ<br>1YCM (NGH) | 20<br>20 |
| 11 | Fibroblast growth factor 1 (FGF1)<br>(*Homo sapiens*) | P05230 | 1RG8<br>1HKN (N2M) | 1.1<br>2.0 | 1DZD<br>1RML (NTS) | 24<br>26 |
| 12 | Peptidyl-prolyl cis-trans isomerase FKBP1A<br>(*Homo sapiens*) | P62942 | 2PPN<br>1J4H (SUB) | 0.9<br>1.8 | 1FKR<br>1F40 (GPI) | 20<br>10 |
| 6 proteins have only holo structures |  |  |  |  |  |  |
| 13 | UDP-3-O-acetyl-N-acetylglucosamine deacetylase<br>(*Aquifex aeolicus*) | O67648 | 3P3C (3P3) | 1.3 | 2JT2 (C90) | 25 |
| 14 | E3 ubiquitin-protein<br>ligase Mdm2<br>(*Homo sapiens*) | Q00987 | 4OGN (2U5) | 1.4 | 2LZG (13Q) | 5 |
| 15 | Dihydrofolate<br>reductase<br>(*Homo sapiens*) | P00374 | 1KMV (LII) | 1.1 | 1YHO (TRR) | 25 |
| 16 | Collagenase 3<br>(*Homo sapiens*) | P45452 | 4L19 (1UA) | 1.7 | 1FM1 (WAY) | 30 |
| 17 | CRE-binding protein<br>(CREBP)<br>(*Homo sapiens*) | Q92793 | 5J0D (6F9) | 1.1 | 2L84 (J28) | 20 |
| 18 | E3 ubiquitin-protein<br>ligase XIAP<br>(*Homo sapiens*) | P98170 | 3HL5 (9Jz) | 1.8 | 1TFT (997) | 1 |

The table lists the 18 selected proteins, categorized by those with only apo structures (7 proteins), those with both apo and holo structures (5 proteins) and those with only holo structures (6 proteins). For each protein, the UniProt ID, PDB IDs of X-ray and NMR structures, X-ray resolution (Å), and the number of available NMR conformers are provided.

**Table 2:**
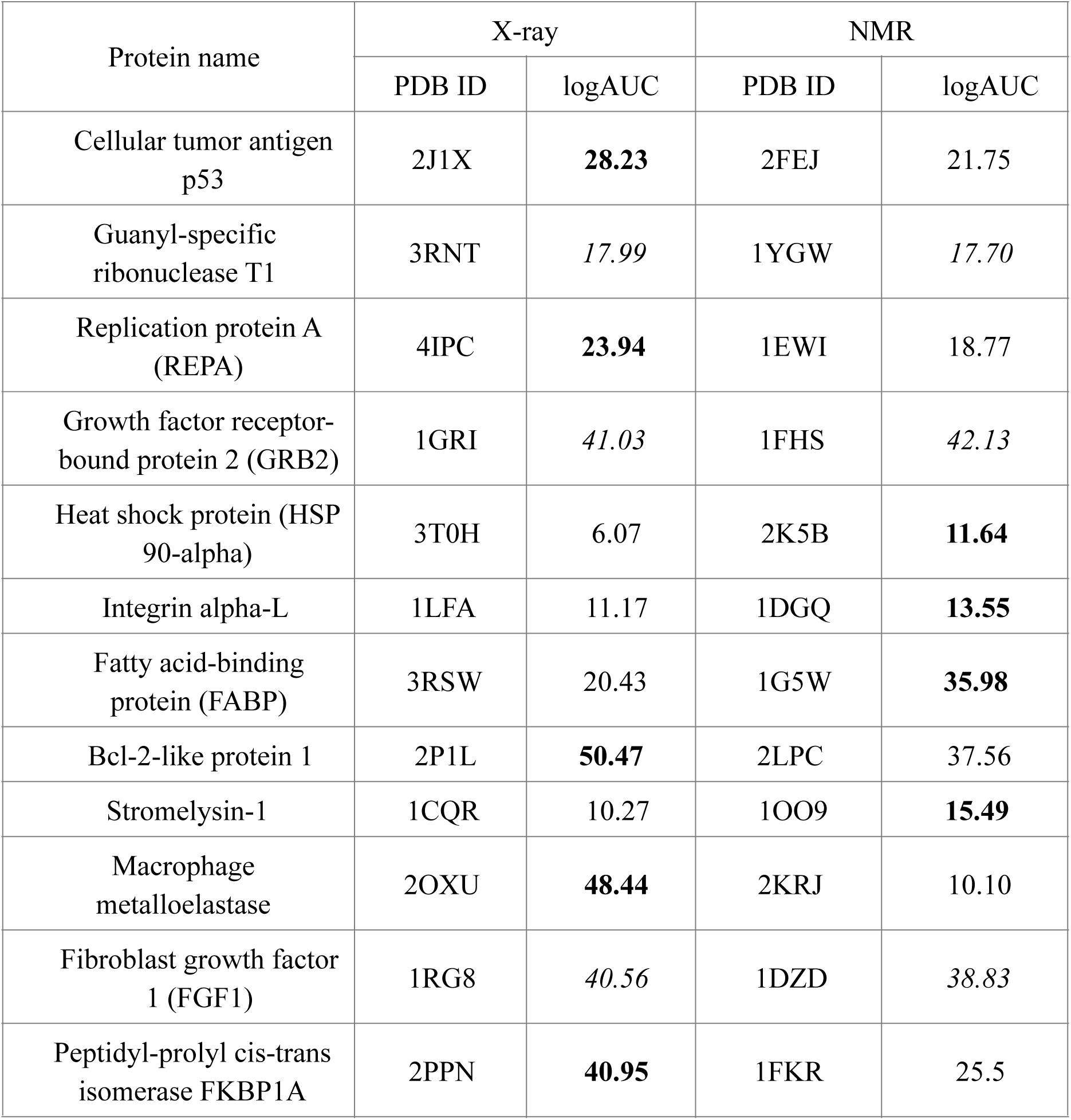
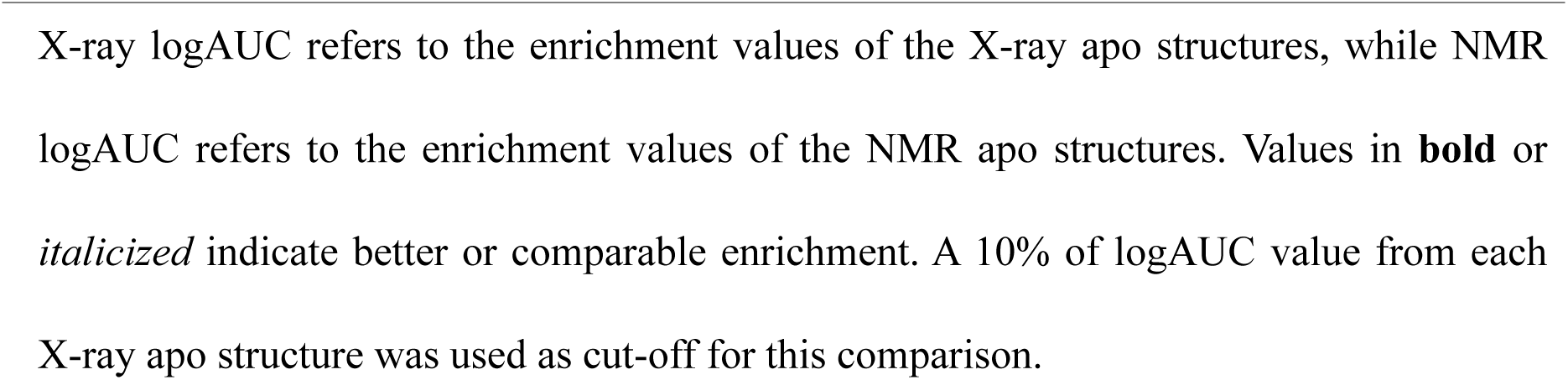
Ligand enrichment of X-ray and NMR apo structures.

The poor virtual screening performance of HSP90, Integrin alpha-L, and Stromelysin-1 apo structures can be attributed to their binding pocket conformation. In the case of HSP90, both X-ray (PDB ID: 3T0H; 246 Å^3^) and NMR (PDB ID: 2K5B; 278 Å^3^) apo structures exhibited significantly smaller binding pocket volumes, compared to the X-ray holo structure of the same protein (PDB ID: 4O0B; 528 Å^3^). The “semi-closed” binding pocket of the apo structures likely restricted their ability to accommodate ligands. When docking screens was conducted against the holo X-ray structure of HSP90 (PDB ID:4O0B), a significant improvement in ligand enrichment (logAUC = 32.41) was observed, supporting this hypothesis. Similarly, the poor performance of Stromelysin-1 and Integrin alpha-L apo structures can also be attributed to their partially closed binding pockets, which restricted ligand accommodation, causing docked ligands to remain predominantly solvent-exposed. When the holo X-ray structures of Stromelysin-1 (PDB ID: 1CIZ) and Integrin alpha-L (PDB ID: 4IXD) were used in virtual screening, improved ligand enrichments were observed, with logAUC values increasing to 15.84 for Stromelysin-1 and to 25.34 for Integrin alpha-L, respectively.

Apo protein structures are determined in the absence of a ligand and often adopt conformations that may not allow ligand binding. As a result, apo structures generally exhibit lower ligand enrichment in virtual screening compared to their respective holo structures. However, when considering only apo structures that performed better that random selection, we observed that X-ray apo structures outperformed NMR apo structures in more than 50% of cases. This trend may be attributed to the larger SASA and greater exposure of polar residues in the binding pockets of X-ray apo structures. On average, X-ray apo structures exhibited a higher total SASA (∼14,032 Å²) compared to NMR apo structures (∼9,617 Å²), suggesting that NMR structures adopt a more compact conformation, which has been previously observed (14-16). Similarly, polar residue exposure was greater in X-ray structures (∼9,972 Å²) than in NMR (∼6,787 Å²), reinforcing the idea of improved pocket accessibility. Greater exposure of polar residues in X-ray apo structures may facilitate hydrogen bonding and electrostatic interactions with docked ligands, enhancing their recognition. Illustrative comparison of these properties is provided in Figure S2.

### 2.3 Ligand enrichment of holo structures

To evaluate the virtual screening performance of holo structures, we docked each library molecule to all X-ray/NMR holo structures, and calculated logAUC values of the enrichment plots for both types of structures (Figure S3). Out of the 11 X-ray/NMR holo structures, ligand enrichment (logAUC) of X-ray holo structures was better, comparable, and worse than that of NMR holo structures in 27%, 64%, and 9% of cases, respectively. The differences in ligand enrichment between X-ray and NMR holo structures were not statistically significant, with a p-value of 0.131.

**Table 3:** Ligand enrichment of single X-ray and NMR holo structures.

| **Table 3: Ligand enrichment of single X-ray and NMR holo structures.** |  |  |  |  |
| --- | --- | --- | --- | --- |
| Protein name | X-ray |  | NMR |  |
|  | PDB ID | X-ray logAUC | PDB ID | NMR logAUC |
| Bcl-2-like protein 1 | 3ZLR | 51.99 | 2O1Y | **58.71** |
| Macrophage metalloelastase | 3F17 | **52.33** | 1YCM | 21.32 |
| Fibroblast growth factor 1 (FGF1) | 1HKN | *36.93* | 1RML | *39.35* |
| Stromelysin-1 | 1CIZ | *15.84* | 2JT6 | *15.87* |
| Peptidyl-prolyl cis-trans isomerase FKBP1A | 1J4H | *32.74* | 1F40 | *30.95* |
| UDP-3-O-acetyl-N-acetylglucosamine deacetylase | 3P3C | *50.22* | 2JT2 | *47.03* |
| E3-ubiquitin ligase MDM2 | 4OGN | *41.07* | 2LZG | *40.47* |
| Dihydrofolate reductase | 1KMV | **57.51** | 1YHO | 24.39 |
| Collagenase 3 | 4L19 | **38.13** | 1FM1 | 27.78 |
| CRE-binding protein (CREBP) | 5J0D | *18.17* | 2L84 | *18.51* |
| E3 ubiquitin ligase XIAP | 3HL5 | *65.32* | 1TFT | *62.24* |

**X-ray logAUC**, enrichment values of X-ray holo structures. **NMR logAUC**, enrichment values of NMR holo structures. Values in **bold** or *italic* indicate better or comparable enrichment. A 10% of logAUC value from each X-ray holo structure was used as cut-off for this comparison.

Unlike apo structures, all holo structures outperformed random selection in ligand enrichment. The presence of a cognate ligand in the target protein structure enhances the definition of the binding pocket, creating a more favourable environment for docking. When comparing single X-ray and NMR holo structures over 11 proteins, the majority (over 60%) exhibited comparable virtual screening performance, suggesting that, the influence of experimental structure type (X-ray vs. NMR) on virtual screening performance is less pronounced in holo structures, than in apo structures.

Interestingly, ligand enrichment was notably low for the Stromelysin-1 protein (logAUC = 15.84 for X-ray holo and logAUC = 15.87 for NMR holo), despite well-defined binding pockets in both X-ray and NMR holo structures. This poor performance is likely attributed to the low chemical similarity (Tc values in the range of 0.03 – 0.36 for both structure types) between ligands in the virtual screening library and the cognate ligand in both structures (Figure S4). Consequently, docked ligands stay largely solvent-exposed, with significantly worse docking scores than that of the cognate ligand (Figure S5).

Next, we compared the virtual screening performance between multiple holo X-ray structures and multiple holo NMR conformers. Among the eight proteins, the ligand enrichment (logAUC) of the best-performing X-ray structure was better, comparable, and worse than that of the best-performing NMR conformer in 63%, 37%, and 0% of cases, respectively. To further explore these differences and minimise biases from individual protein structures, we conducted pairwise comparisons between all three X-ray structures and all three NMR conformers for each protein, in total 72 comparisons across all eight proteins. This analysis revealed a statistically significant difference in ligand enrichment favoring X-ray holo structures, with p-value of 0.001 (Table 4). Consensus enrichment was also calculated for each protein, by combining docking results of multiple structures (X-ray) and conformers (NMR). For each docked compound (ligands and decoys), the best docking score across all structures/conformers was used for the ranking of that compound. Our results showed that consensus enrichment of multiple holo X-ray structures was better, comparable, and worse than that of multiple holo NMR conformers in 63%, 25%, and 12% of proteins, respectively (Table 4, Figure S6). Moreover, for X-ray holo structures, consensus enrichment outperformed the best performing X-ray holo structure in 12.5% of protein cases, the second-best X-ray holo structure in 62.5% of protein cases, and the third-best X-ray holo structure in all protein cases (100%). Similarly, for NMR conformers, consensus enrichment outperformed the best performing single NMR conformer in 12.5% of protein cases, the second-best conformer in 50% of cases and the third-best conformer in 75% of cases. These findings suggest that incorporating multiple structures/conformers in docking screens can improve performance by accounting for variations in protein-ligand interactions, reducing the risk of relying on single preselected structure. Moreover, in real-world applications where the true ligands are unknown, it is difficult to identify the best-performing structure a priori. Hence, selecting multiple X-ray structures with chemically diverse cognate ligands (rather than diverse conformations from NMR ensemble) may capture different ligand binding motifs, potentially improving ligand ranking in molecular docking.

**Table 4:**
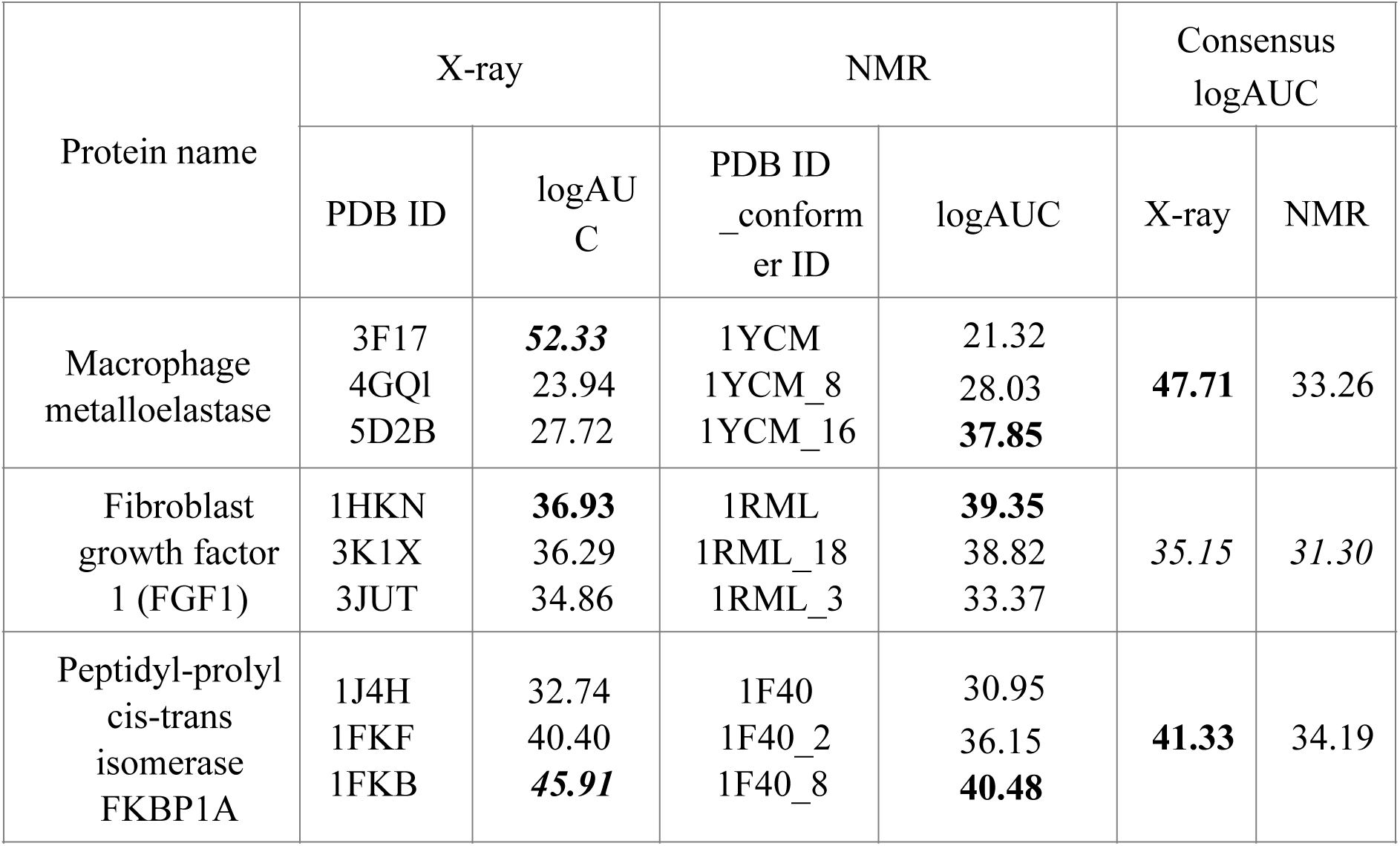
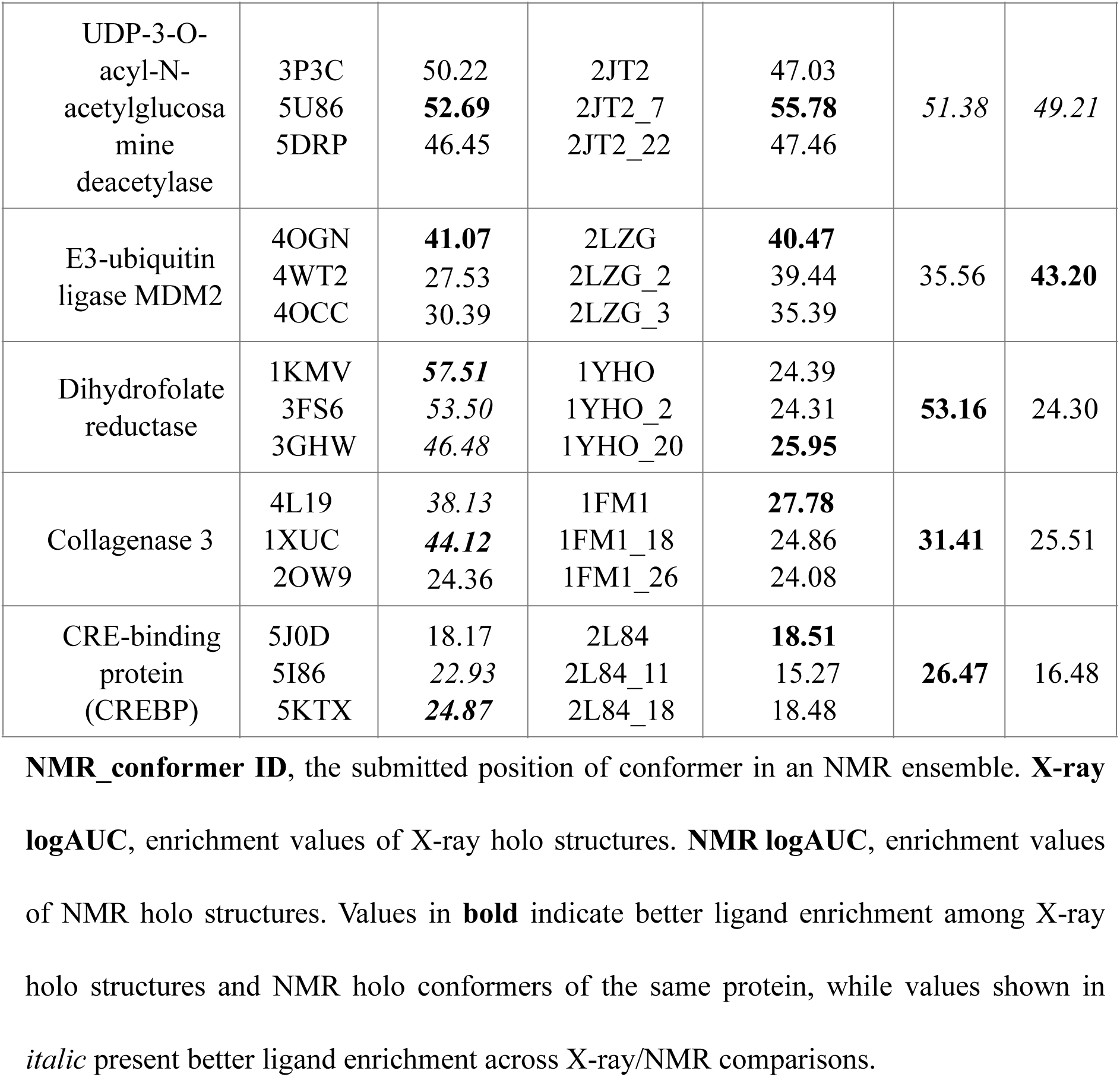
Ligand enrichment of multiple X-ray and NMR holo structures/conformers.

### 2.4 Analysis

Next, we conducted a comprehensive analysis of ligand properties, protein features, and protein-ligand interactions to evaluate their impacts on virtual screening performance. We first examined structural deviations in the binding pocket, followed by an assessment of common protein binding site features (volume and SASA, hydrophobicity and the proportion of polar atoms) to determine their correlations with molecular docking outcomes (Table S2). Additionally, to estimate ligand recognition biases during molecular docking, we assessed the chemical similarity between cognate and database ligands. Finally, we analysed protein-ligand interactions, focusing on hydrophobic contacts and hydrogen bonds, to understand their roles in ligand enrichment.

#### 2.4.1 Protein feature assessment

##### 2.4.1.1 Binding site structural deviation

To examine the impact of binding site structural deviations on virtual screening performance, we calculated pairwise RMSD values for the heavy atoms of binding site residues across holo X-ray/NMR structures/conformers for each protein (Table 5). The binding site for each protein was defined as the union of all residues within 6Å distance from the cognate ligands across all X-ray/NMR structures/conformers associated with that protein. Our analysis showed that NMR holo conformers exhibited greater structural diversity (higher RMSD values) than X-ray holo structures in 7 out of 8 proteins (88%). This difference was statistically significant, with p-value of 0.039. However, for one protein (FKBP1A), the RMSD values among NMR conformers were 0.0 Å, indicating negligible structural differences. Despite the lack of structural differences among FKBP1A NMR holo conformers (RMSD = 0.0 Å for binding site residues), differences were observed in cognate ligand binding modes. Specifically, after superimposing the 2^nd^ and 8^th^ conformer in the deposited NMR ensemble onto the first deposited conformer, we calculated the RMSD of the heavy atoms between their cognate ligands and the cognate ligand of the first conformer. This result revealed RMSD of 3.28 Å and 4.15 Å, respectively, indicating substantial differences in ligand binding modes among these three NMR holo conformers.

**Table 5:**
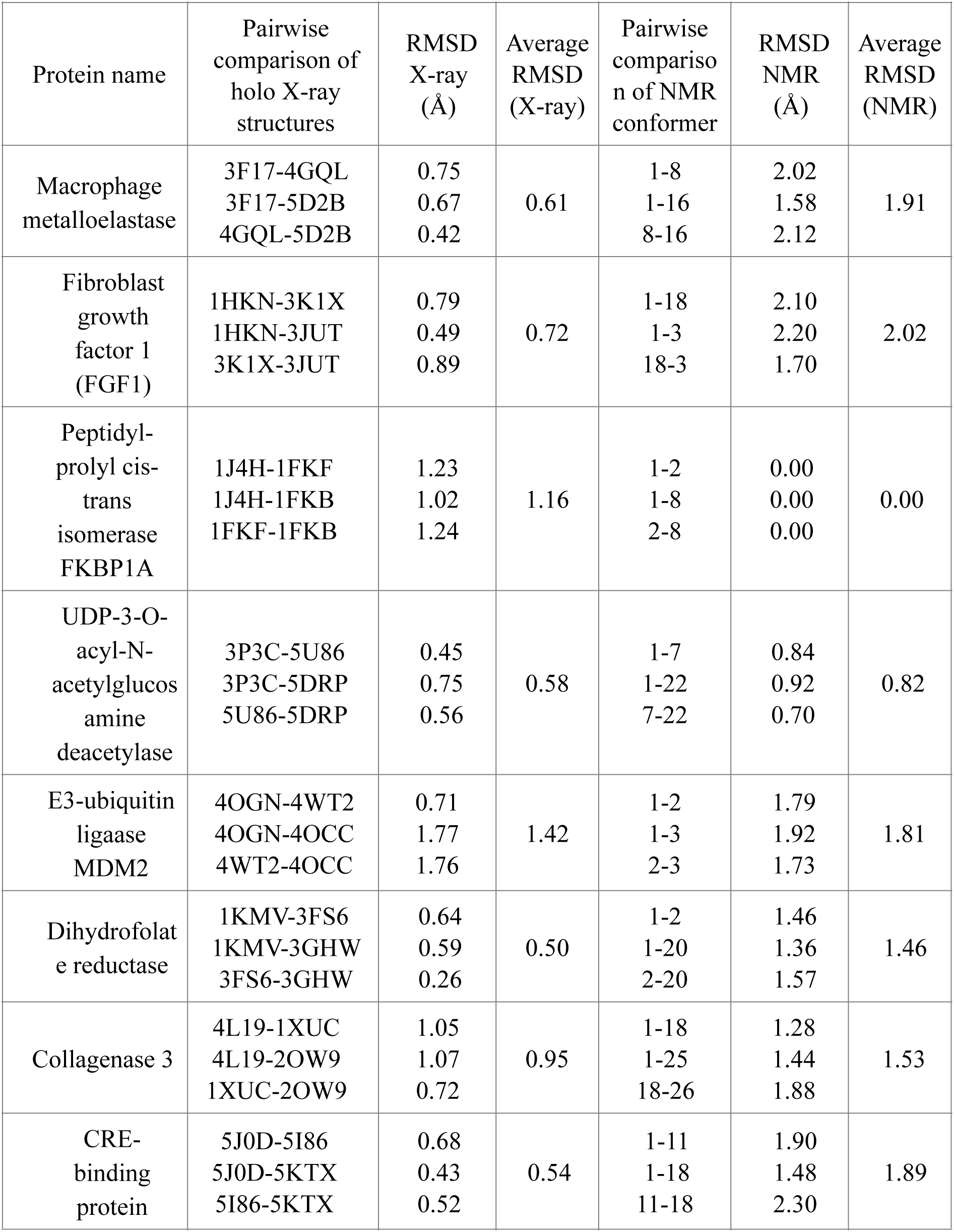
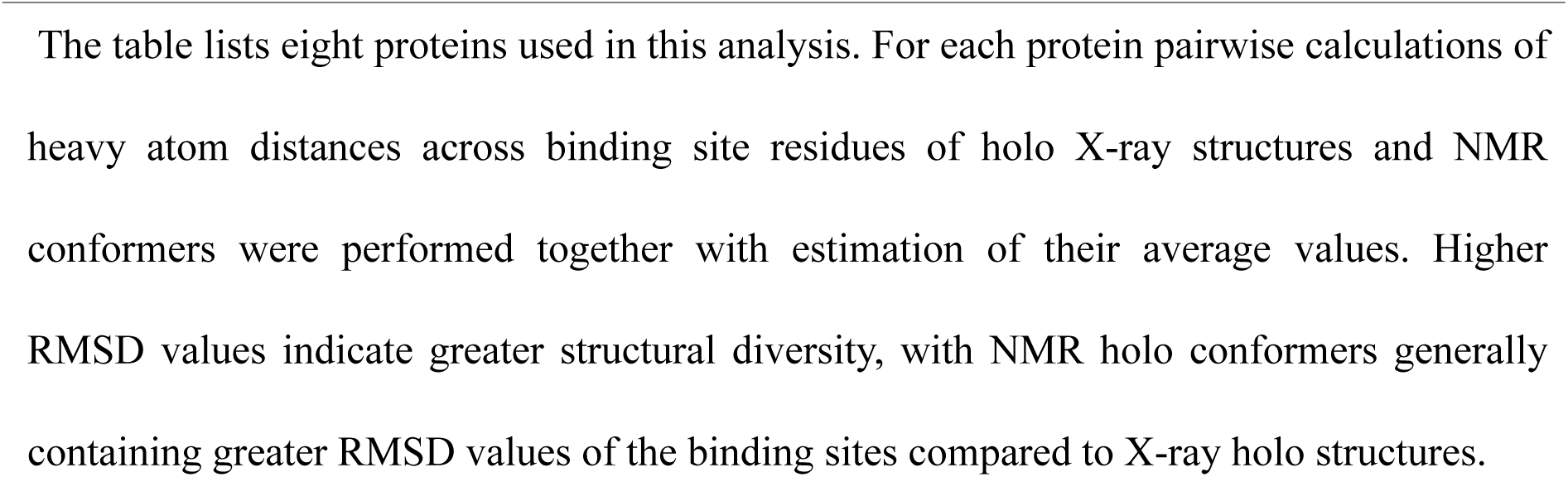
Binding site structural deviations of multiple holo X-ray structures and NMR conformers.

Our results suggest that the binding site conformation in different NMR conformers exhibit greater structural diversity compared to that in different X-ray holo structures. This aligns with findings from Schneider et al. (17), which highlighted that NMR structures tend to display higher conformational heterogeneity and greater uncertainties in backbone and side-chain conformations compared to X-ray structures, which can impact their performance as structural templates for protein design. As shown in our previous comparison, X-ray holo structures outperformed NMR holo conformers, despite the greater binding site diversity observed in NMR conformers (Table 4). This finding suggests that simply increasing structural diversity of the binding site does not necessarily improve virtual screening performance. Instead, selecting multiple holo structures with chemically diverse cognate ligands, may be a more effective strategy to improve ligand recognition performance.

##### 2.4.1.2 Binding site feature assessment

Previously, we demonstrated that binding site volume can impact virtual screening performance, as evidenced with the X-ray apo structure of HSP90 protein. To further explore this, we analysed whether differences in binding site features (volume, total, polar, apolar SASA, hydrophobicity and proportion of polar atoms) between X-ray structures and NMR holo conformers correlate with differences observed in ligand enrichment. The analysis revealed weak correlations across all features, with binding site volume showing an R² of 0.05 and total, polar, and apolar SASA differences yielding R² values of 0.04, 0.12, and 0.00, respectively. Hydrophobicity and the proportion of polar atoms showed no correlation (R² = 0.00). Full results are provided in Figures S7 and S8.

#### 2.4.2 Chemical similarity between cognate and database ligands

In this study, X-ray and NMR holo structures of the same protein were solved in complex with different cognate ligands. The binding mode of the cognate ligand influences the shape of the binding site and, consequently, ligand recognition in molecular docking. Therefore, for all eight proteins, we calculated and compared the chemical similarity distributions between their cognate ligands of X-ray/NMR structures and the 10 database ligands used in molecular docking (Figure S9). Each protein had three distinct X-ray holo structures and one NMR holo structure, resulting in 240 total comparisons for X-ray (3 cognate ligand x 8 proteins x 10 database ligands), and 80 for NMR (1 cognate ligand x 8 proteins x 10 database ligands). Among the X-ray cognate ligands, only two – from macrophage metalloelastase (PDB ID: 5D2B) and CRE-binding protein (PDB ID: 5KTX) – showed high chemical similarity (Tc > 0.7) to one database ligand. Similarly, among NMR cognate ligands, two – from MDM2 (PDB ID: 2LZG) and DHFR (PDB ID: 1YHO) – showed Tc > 0.7 to one database ligand. These results suggest that, overall, the majority of cognate ligands from X-ray and NMR holo structures are chemically distinct from database ligands used in molecular docking.

#### 2.4.3 Protein-ligand interactions feature assessment

Ligand enrichment performance is directly influenced by the nature of protein-ligand interactions in docking poses. To assess whether differences in virtual screening performance correlated with differences in specific types of protein-ligand interactions, we analysed hydrophobic C-C contacts and hydrogen bonds formed between protein binding site residues and docked ligands. The differences in the numbers of hydrophobic contacts formed at different distances (4 Å/5 Å/6 Å) present higher correlation with the differences in the ligand enrichment than binding site features, with *R*^2^ values of 0.16, 0.28, and 0.35, respectively (Figure 2).

**Figure 2.**
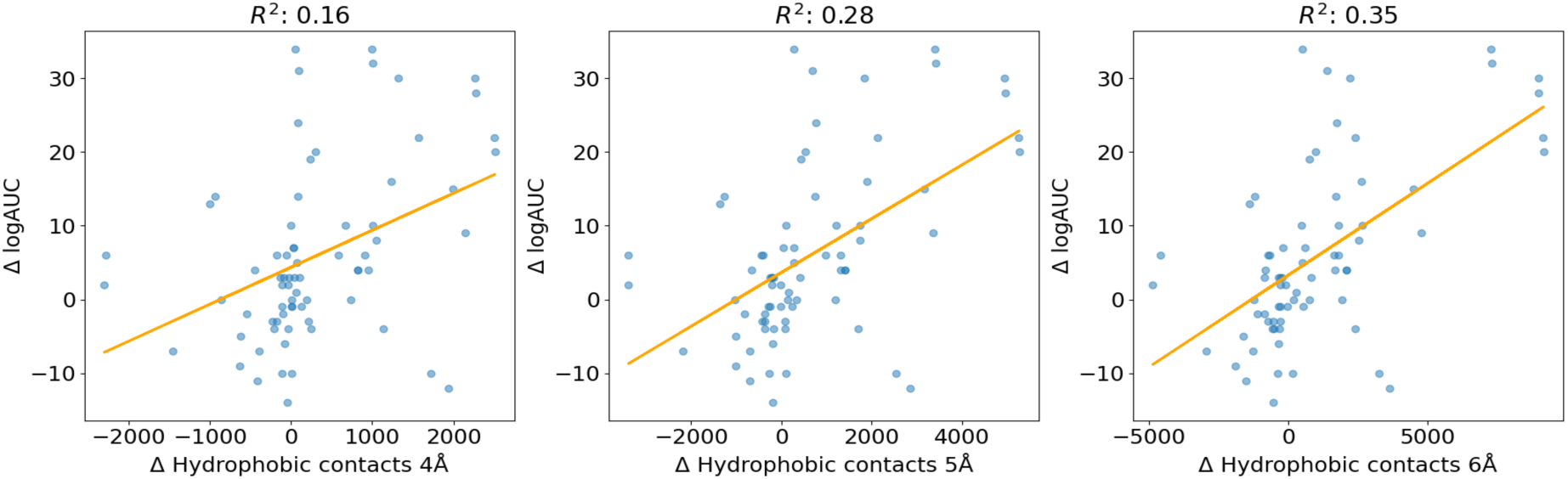
Scatter plot of the X-ray/NMR regimes. The plot illustrates the correlation between the differences in the ligand enrichment of the multiple X-ray/NMR holo structures/ conformers (ΔlogAUC) and the differences in the number of hydrophobic contacts formed between protein residues and top-scored docked ligand at distances of 4 Å; 5 Å and 6 Å (ΔHydrophobic contacts).

Similarly, differences in hydrogen bond interactions showed the highest correlation with the differences in ligand enrichment (R^2^ = 0.43, Figure 3). These findings indicate that differences in both hydrophobic contacts (6 Å) and hydrogen bonding interactions significantly contribute to the differences in ligand enrichment between X-ray and NMR holo structures, with hydrogen bonding being the most important feature. This trend aligns with the fundamental principles of docking energy functions, such as the one used by Glide, which contains hydrogen bonding and hydrophobic interactions in the energy calculations.

**Figure 3:**
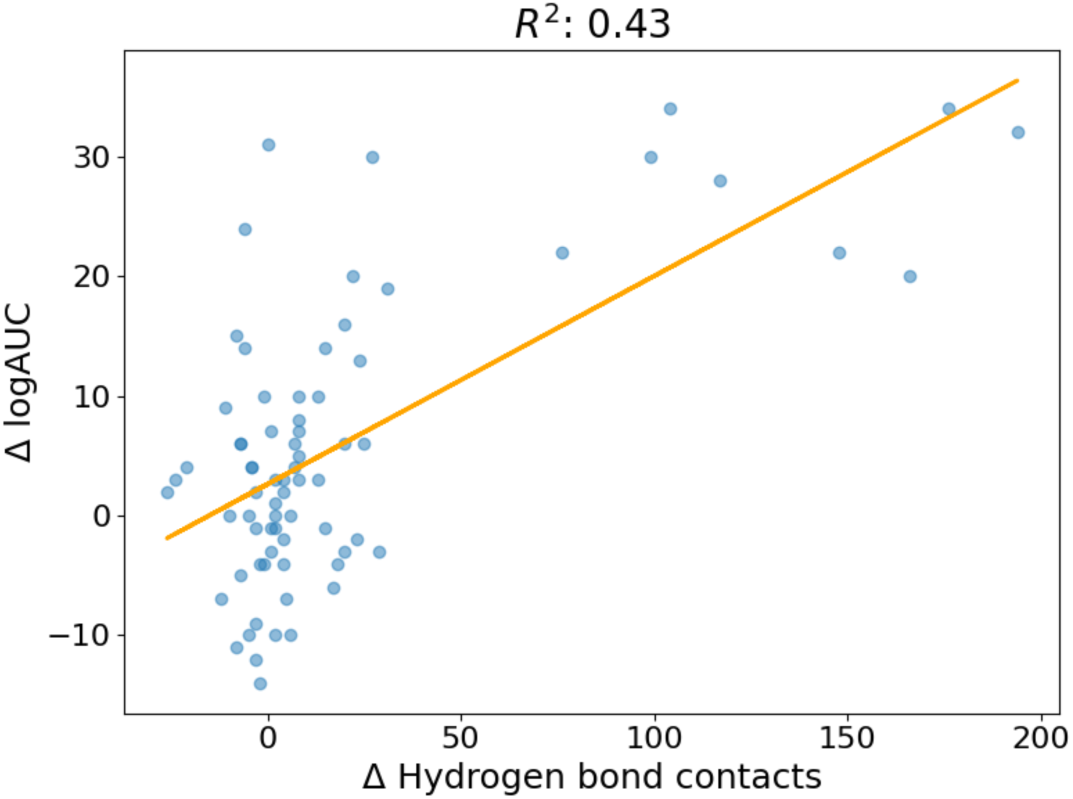
Scatter plot of the X-ray/NMR regimes. The plot illustrates the correlation between the differences in the ligand enrichment of the multiple X-ray/NMR holo structures/ conformers (ΔlogAUC) and the differences in the number of hydrogen bond contacts formed between protein residues and top-scored docked ligand (ΔHydrogen bond contacts).

## 3. Conclusions

This study systematically compared the virtual screening performance of X-ray and NMR holo structures/conformers. Our findings indicate that X-ray holo structures generally outperform NMR holo conformers, with differences in hydrogen bonding and hydrophobic contacts being the most important features for virtual screening performance. While NMR conformers in the same ensemble exhibited greater structural diversity of the binding sites, in comparison to multiple X-ray holo structures of the same protein; this did not translate into improved virtual screening performance, reinforcing the idea that structure selection should prioritise those structures with chemically diverse cognate ligands rather than considering sheer conformational heterogeneity of the binding sites.

Our study also has limitations. The correlation analysis was performed on a limited dataset (eight proteins), and trends may vary with larger datasets or different protein families. Additionally, our findings are specific to the Glide docking algorithm and OPLS3e force field—while these methods are widely used in structure-based drug design, different scoring functions (e.g., DOCK4, AutoDock, GOLD) may weigh interaction terms differently. Future work could explore whether machine learning models trained on protein and protein-ligand interaction features can predict the most suitable protein structure for docking screens. Finally, the role of explicit water molecules in mediating key interactions also remains an open question and could be investigated further by incorporating solvent modeling approaches in docking simulations.

## 4. Materials and Methods

### 4.1 Protein target selection

Protein targets in this study were selected from the UniProt database (18) and the Protein Data Bank (PDB) (19), following five criteria: 1) only proteins with both X-ray and NMR holo and apo structural data available were included; 2) proteins were required to contain more than 90 amino acids in a single chain; 3) among holo structures, we selected those with a buried ligand, defined as having at least one pair of non-hydrogen atoms from the protein and ligand within a distance cut-off of 4 Å; 4) the selected X-ray structures had to meet a resolution cutoff of 3.0 Å or better, prioritising the best resolution available; and 5) solid-state NMR structures, unpublished structures, and those with missing residues or with mutations in the binding pocket were excluded. Ultimately, eighteen proteins meeting these criteria were selected for this study (Table 1).

The sample of 18 proteins can be divided into two functional classes based on their biological roles: Class I includes 11 proteins that act as catalysts of biological reactions. This class includes hydrolases (HSP90, UDP deacetylase, Collagenase 3, Stromelysin-1, Metalloelastase, Guanyl-specific Ribonuclease T1), transferases (CREBP), isomerases (FKBP1A), ligases (E3-ubiquitin ligase MDM2, E3-protein ligase XIAP), and oxidoreductase (DHFR). Class II includes seven proteins that are not enzymes. This class consists of regulatory proteins (P53, Bcl-2, FGF1, and RepA), intracellular transporter of long-chain fatty acids (FABP), adaptor protein (GRB2) that presents an essential link between GFR and RAS signalling pathway, and the cell adhesion receptor (Integrin alpha) that plays an important role in various immune processes. Most of the proteins are also interesting targets for drug discovery such as P53, E3-ubiquitin ligase MDM2, E3-ubiquitin ligase XIAP, Bcl-2, and HSP90 (20–24).

To perform comparisons of virtual screening performance, the 3D structures of these 18 proteins determined in apo and/or holo state, were collected. Among them, 12 proteins have one apo X-ray structure and one apo NMR structure, and 11 proteins have at least one holo X-ray structure and one holo NMR structure (Table S3). For each of these 11 proteins, the X-ray structure with best resolution was selected, together with the first conformer of the NMR ensemble deposited in PDB.

### 4.2 Multiple X-ray structures and NMR conformers

Among the 11 proteins with at least one holo X-ray structure and one holo NMR structure, 8 proteins have at least three holo X-ray structures (Table S4). For each of these 8 proteins, the holo NMR structure typically contains more than ten conformers, except for that of E3-ubiquitin ligase MDM2 (PDB ID: 2LZG, five conformers). For each of the eight proteins, we employed a two-step selection process to identify two additional X-ray holo structures beyond the structure that was already selected with the best resolution: 1) we first filtered all available X-ray holo structures, selecting those whose cognate ligand is not chemically similar (Tanimoto coefficient (Tc) < 0.7) to that in the first structure; 2) from these selected structures, we picked two additional structures with the second and third best resolutions. To match the three X-ray structures, for each of these 8 proteins, two more conformers were selected from the same NMR ensemble using the following algorithm: 1) the 2^nd^ conformer has the maximal binding site (defined as residues within 6Å of the ligand) heavy atom RMSD from the 1st conformer; 2) the 3rd conformer has the maximal binding site heavy atom RMSD from the 1^st^ and 2^nd^ conformer.

### 4.3 Chemical library used in docking screens

To evaluate the cognate docking and virtual screening performance of X-ray and NMR structures, a chemical library was prepared for each protein, consisting of: 1) cognate ligands that were solved in complex with the X-ray and/or NMR structures; 2) 10 known ligands retrieved from BindingDB (25) (Table S5); and 3) 50 property-matching decoys per known ligand, selected from the ZINC database (26) using DUD-E protocol (27). The property-matching decoys were selected for their chemical dissimilarity to the known ligands (Tc < 0.4) and were considered non-binders.

### 4.4 Molecular docking – Method description

All protein structures were pre-processed using the Preparation Wizard module (Protein Preparation Wizard, Schrödinger, LLC, New York, NY). The 3D structures of the chemical library molecules were prepared with LigPrep (LigPrep, Schrödinger, LLC, New York, NY), while molecular docking was performed on both holo and apo structures using the Glide SP module (Glide, Schrödinger, LLC, New York, NY) (28).

For the holo structures, a receptor grid was generated using the Receptor Grid Generation Panel within the Glide Suite. A cubic grid box, defined as the inner box (15 Å per side), and a larger cubic box, designated as the outer box (20 Å per side), were centred on the centroid of the crystal ligand. In the case of the apo structures, the receptor grid was generated for the same binding pocket as determined in the corresponding holo structure. This was accomplished by superposing the holo and apo structures to ensure accurate localisation of the binding site. During the docking procedure, all cofactors and metal ions present in the binding pocket were retained to maintain the integrity of the active site. The OPLS3e force field was employed for identifying and ranking the docking poses (29).

### 4.5 Virtual screening performance evaluation

In this study, virtual screening performance was assessed by ligand enrichment using the logAUC metric (8, 9). The difference in ligand enrichment yielded using X-ray and NMR structures was assessed in two ways: 1) The difference was considered significant if it exceeded 10% of the logAUC value of the X-ray structure (arbitrary approach); 2) The distribution of differences was tested for normality using the Shapiro-Wilk test (30). If the differences were normally distributed at a 95% confidence level, single-factor ANOVA analysis was used (31). Otherwise, the Wilcoxon signed-rank test (32) was conducted to estimate the statistical significance of the differences using the R package (33).

### 4.6 Feature assessment

To identify features contributing to differences in ligand enrichment, several analyses were performed. First, 2D fingerprints (34) of cognate ligands in experimental structures and database ligands used in virtual screening — were generated and compared using OpenBabel (35). This comparison was used to quantify chemical similarity (Tc values) between the two ligand sets. Second, percentages of ligand pairs that fall within 0.0 – 0.7 Tc range (low to moderate similarity) and the 0.7 – 1.0 range (high similarity) were estimated. This allowed us to assess whether ligands in NMR or X-ray structures exhibited different trends of chemical similarity against database ligands, providing insights into potential artefacts in ligand enrichment results. Third, violin plots were generated to visualize chemical similarity distributions for X-ray and NMR structures, using kernel density estimation (KDE), which provides a smoothed representation of probability density function for Tc values. Additionally, three further analyses were conducted to explore correlation between ligand enrichment differences and differences in protein/ligand features:

1. Fpocket (36) was used to calculate binding site volume, polar/apolar/total SASA, hydrophobicity, and the proportion of the polar atoms in the binding pocket.
2. PLIP (37) was used to assess the number of hydrogen bonds formed between X-ray/ NMR structures/conformers binding site residues and best-scored ligand docking poses.
3. The number of C-C hydrophobic contacts (at distances of 4Å, 5Å, and 6Å) formed between X-ray/NMR structure/conformers binding site residues and best-scored ligand docking poses.

The differences in protein/ligand features (e.g. ΔVolume, ΔSASA, Δhydrophobicity, and the Δproportion of the polar atoms) and ligand enrichment values (ΔlogAUC) between X-ray and NMR structures for each protein were calculated by subtracting the values derived from NMR structures from those obtained from X-ray structures. Linear regression analysis was then performed on these differences to assess their correlation, with R^2^ values being calculated. As mentioned previously, eight proteins contained three X-ray holo structures and three NMR holo conformers. To eliminate biases, differences were calculated and compared across all X-ray structures and all NMR conformers for each protein. Pairwise comparison was conducted between all X-ray structures and NMR conformers, resulting in a 3x3 matrix per protein.

## Supplementary Material Description

Supplementary Material includes five tables and nine figures supporting the main manuscript. Table S1 summarizes the accuracy of cognate ligand docking poses across X-ray and NMR structures. Table S2 presents key binding pocket and protein-ligand interaction features, including SASA, hydrophobic contacts, hydrogen bonds, and pocket volumes. Tables S3-S5 provide details on protein subsets, structural identifiers, and chemical libraries used in virtual screening. Figures S1-S9 display enrichment curves, structural comparisons, chemical similarity distributions, docking pose visualisations, and correlations between structural features and virtual screening performance.

## Author Contributions

Srdan Masirevic performed all molecular docking simulations, analyses, and wrote the manuscript. Dr. Hao Fan supervised the research, provided conceptual guidance, and reviewed the manuscript.

## Acknowledgements

This research was supported by SINGA scholarship and Bioinformatics Institute at A*STAR. I am sincerely thankful to Dr. Hao Fan, Dr. Chandra Verma from Bioinformatics Institute at A*STAR and Dr. Jayaraman Sivaraman from Department of Biological Sciences at National University of Singapore, for their guidance and insightful discussions throughout this project.

