## Supplementary Information for "Comparative study of protein X-ray and NMR structures: molecular docking-based virtual screening"

Virtual screening, Molecular docking, Protein-ligand interactions, X-ray crystallography, NMR spectroscopy.

| **Table S1: Cognate ligand geometry assessment.** | | | | |
| --- | --- | --- | --- | --- |
| Protein  name | PDB ID | | X-ray -  RMSD (Å) of the top-scored pose (lowest RMSD) | NMR -  RMSD (Å) of the top-scored pose (lowest RMSD) |
|  | X-ray  (best resolution) | NMR |  |  |
| Bcl-2-like protein 1 | 3ZLR | 2O1Y | **0.30 (0.30)** | 4.15 (4.15) |
| Stromelysin-1 | 1CIZ | 2JT6 | 4.53 (4.53) | **1.41 (1.41)** |
| Macrophage metalloelastase | 3F17 | 1YCM | 3.06 (**2.27**) | **1.35 (1.34)** |
| Fibroblast growth factor 1 (FGF1) | 1HKN | 1RML | **2.46 (2.46)** | 3.95 (3.95) |
| Peptidyl-prolyl cis-trans isomerase FKBP1A | 1J4H | 1F40 | **2.97 (2.97)** | **2.61 (2.61)** |
| UDP-3-O-acyl-N-acetylglucosamine deacetylase | 3P3C | 2JT2 | **0.24 (0.24)** | **1.01 (0.87)** |
| E3-ubiquitin ligase MDM2 | 4OGN | 2LZG | **0.50 (0.50)** | **1.62 (0.85)** |
| Dihydrofolate reductase | 1KMV | 1YHO | **0.19 (0.19)** | **2.02 (2.02)** |
| Collagenase 3 | 4L19 | 1FM1 | **0.30 (0.30)** | 5.09 (4.72) |
| CRE-binding protein (CREBP) | 5J0D | 2L84 | **2.73 (2.59)** | **1.92 (1.92)** |
| E3 ubiquitin ligase XIAP | 3HL5 | 1TFT | **0.63 (0.63)** | **0.89 (0.89)** |
| The table lists 11 proteins used for this analysis. PDB codes of X-ray (best resolution) and NMR structures, together with RMSD values for ligand docking poses with best docking energy (top-scored), as well as ligand poses with lowest RMSD value among all the docking poses sampled (shown in brackets), are listed. **Bold** indicates that we reproduce cognate ligand conformation accurately (RMSD < 3 Å), with docking. | | | | |

| **Table S2: Protein binding pocket and protein-ligand interaction feature results.** | | | | | | | | | | | | | | | | | | | | | | |
| --- | --- | --- | --- | --- | --- | --- | --- | --- | --- | --- | --- | --- | --- | --- | --- | --- | --- | --- | --- | --- | --- | --- |
| Protein name | X-ray logAUC | NMR logAUC | Hydrophobic contacts | | | | | | Hydrogen bonds  X-ray/NMR | | Binding pocket volume (Å ^3^) | | Total SASA  X-ray/NMR  (Å ^2^) | | Polar SASA  X-ray/NMR  (Å ^2^) | | Apolar SASA  X-ray/NMR  (Å ^2^) | | Hydrophobicity  X-ray/NMR  (Å ^2^) | | Polar atom proportion  X-ray/NMR | |
|  |  |  | 4Å  X-ray/NMR | | 5Å  X-ray/NMR | | 6Å  X-ray/NMR | |  |  |  |  |  |  |  |  |  |  |  |  |  |  |
| Macrophage metalloelastase | 52 | 21 | 230 | 133 | 1543 | 862 | 3694 | 2300 | 18 | 18 | 750 | 375 | 236 | 98 | 85 | 25 | 151 | 73 | 32 | 38 | 42 | 42 |
|  | 24 | 28 | 107 | 146 | 608 | 773 | 1441 | 1944 | 22 | 24 | 802 | 1044 | 194 | 300 | 64 | 124 | 130 | 176 | 33 | 76 | 42 | 75 |
|  | 28 | 38 | 158 | 152 | 907 | 799 | 2123 | 1966 | 19 | 24 | 962 | 828 | 268 | 232 | 76 | 89 | 192 | 143 | 35 | 56 | 37 | 85 |
| Fibroblast growth factor 1 (FGF1) | 37 | 39 | 58 | 155 | 374 | 743 | 931 | 1777 | 50 | 46 | 265 | 511 | 36 | 112 | 13 | 42 | 23 | 70 | -13 | 21 | 39 | 40 |
|  | 36 | 39 | 369 | 602 | 829 | 1200 | 1506 | 2027 | 47 | 27 | 261 | 263 | 85 | 64 | 61 | 16 | 24 | 48 | -9 | 16 | 56 | 29 |
|  | 35 | 33 | 397 | 506 | 833 | 1045 | 1454 | 1736 | 45 | 71 | 0 | 0 | 0 | 0 | 0 | 0 | 0 | 0 | 0 | 0 | 0 | 0 |
| Peptidyl-prolyl cis-trans isomerase FKBP1A | 33 | 31 | 266 | 307 | 1298 | 1314 | 3088 | 3173 | 28 | 24 | 455 | 500 | 154 | 112 | 78 | 29 | 76 | 83 | 131 | 63 | 52 | 26 |
|  | 40 | 36 | 2454 | 1630 | 4681 | 3268 | 7961 | 5860 | 13 | 17 | 528 | 495 | 193 | 118 | 92 | 32 | 100 | 86 | 114 | 63 | 59 | 26 |
|  | 46 | 40 | 2303 | 1724 | 4473 | 3490 | 7680 | 6017 | 16 | 23 | 627 | 527 | 148 | 144 | 59 | 46 | 89 | 98 | 55 | 131 | 21 | 58 |
| UDP-3-O-acyl-N-acetylglucosamine deacetylase | 50 | 47 | 214 | 176 | 1080 | 1289 | 2685 | 3022 | 53 | 45 | 1925 | 1289 | 548 | 233 | 261 | 143 | 286 | 90 | 52 | 25 | 77 | 38 |
|  | 53 | 56 | 123 | 296 | 839 | 1273 | 2303 | 3040 | 65 | 36 | 1618 | 1524 | 473 | 382 | 225 | 163 | 248 | 219 | 40 | 19 | 86 | 37 |
|  | 46 | 47 | 185 | 302 | 1006 | 1251 | 2674 | 2962 | 42 | 40 | 1167 | 1050 | 320 | 255 | 144 | 103 | 177 | 152 | 22 | 26 | 38 | 35 |
| E3 ubiquitin ligase MDM2 | 41 | 40 | 313 | 256 | 1395 | 1227 | 3314 | 3020 | 11 | 9 | 783 | 489 | 148 | 106 | 32 | 34 | 116 | 72 | 82 | 63 | 54 | 28 |
|  | 28 | 39 | 2201 | 2616 | 4086 | 4782 | 6659 | 8177 | 6 | 14 | 513 | 541 | 145 | 168 | 29 | 16 | 116 | 152 | 59 | 117 | 29 | 53 |
|  | 30 | 35 | 1979 | 2598 | 3769 | 4787 | 6295 | 7909 | 11 | 18 | 689 | 726 | 217 | 237 | 47 | 33 | 171 | 204 | 131 | 60 | 54 | 32 |
| Dihydrofolate reductase | 58 | 24 | 1147 | 1092 | 4035 | 3750 | 8739 | 8219 | 246 | 142 | 1269 | 933 | 365 | 265 | 118 | 42 | 246 | 223 | 21 | 24 | 31 | 27 |
|  | 54 | 24 | 2418 | 151 | 5585 | 640 | 10437 | 1453 | 169 | 70 | 1074 | 917 | 354 | 265 | 108 | 42 | 246 | 223 | 20 | 24 | 29 | 27 |
|  | 46 | 26 | 2658 | 145 | 5889 | 616 | 10614 | 1438 | 218 | 52 | 1120 | 1210 | 374 | 365 | 121 | 68 | 253 | 297 | 18 | 37 | 32 | 29 |
| Collagenase 3 | 38 | 28 | 199 | 198 | 1306 | 1204 | 3323 | 2850 | 38 | 25 | 1119 | 1152 | 306 | 268 | 145 | 78 | 161 | 190 | 36 | 37 | 43 | 40 |
|  | 44 | 25 | 1435 | 1202 | 3096 | 2665 | 5487 | 4716 | 45 | 14 | 1206 | 933 | 345 | 267 | 172 | 69 | 172 | 198 | 29 | 78 | 42 | 76 |
|  | 24 | 24 | 1329 | 1132 | 2908 | 2564 | 5263 | 4515 | 29 | 23 | 1077 | 956 | 350 | 270 | 179 | 74 | 172 | 196 | 28 | 68 | 43 | 78 |
| CRE-binding protein (CREBP) | 18 | 19 | 170 | 159 | 875 | 883 | 2052 | 2083 | 16 | 15 | 848 | 838 | 214 | 150 | 55 | 78 | 159 | 72 | 34 | 36 | 30 | 40 |
|  | 23 | 15 | 1109 | 63 | 2192 | 455 | 3777 | 1229 | 22 | 14 | 816 | 812 | 293 | 223 | 90 | 89 | 203 | 134 | 127 | 32 | 100 | 35 |
|  | 25 | 18 | 1065 | 1034 | 2188 | 1913 | 3885 | 3272 | 22 | 14 | 669 | 649 | 214 | 164 | 40 | 56 | 174 | 107 | 34 | 42 | 33 | 32 |
| The table lists eight proteins used for multiple holo X-ray vs multiple holo NMR comparison, together with corresponding logAUC values. Hydrophobic contacts, the number of hydrophobic interactions determined between carbon atoms of the best scored docked ligands and carbon atoms of X-ray/NMR binding site atoms, for distances of 4 Å, 5 Å, and 6 Å. Hydrogen bonds, the number of hydrogen bonds formed between X-ray/NMR binding site atoms and top scored ligands. Binding pocket volume, Total/Polar/Apolar SASA, Hydrophobicity and Polar atom proportion are binding pocket features calculated with Fpocket. | | | | | | | | | | | | | | | | | | | | | | |

| **Table S3: Two subsets of the protein database used in docking screens.** | |
| --- | --- |
| X-ray apo vs NMR apo | X-ray holo vs NMR holo |
| Cellular tumor antigen p53 | Bcl-2-like protein 1 |
| Guanyl-specific ribonuclease T1 | Stromelysin-1 |
| Replication protein A (REPA) | *Macrophage metalloelastase* |
| Growth factor receptor-bound protein 2 (GRB2) | *Fibroblast growth factor 1 (FGF1)* |
| Heat shock protein (HSP 90-alpha) | *Peptidyl-prolyl cis-trans isomerase FKBP1A* |
| Integrin alpha-L | *UDP-3-O-acyl-N-acetylglucosamine deacetylase* |
| Fatty acid-binding protein (FABP) | *E3-ubiquitin ligase MDM2* |
| Bcl-2-like protein 1 | *Dihydrofolate reductase* |
| Stromelysin-1 | *Collagenase 3* |
| Macrophage metalloelastase | *CRE-binding protein (CREBP)* |
| Fibroblast growth factor 1 (FGF1) | E3 ubiquitin-protein ligase XIAP |
| Peptidyl-prolyl cis-trans isomerase FKBP1A |  |
| The left column lists proteins used for comparison between X-ray and NMR apo structures, while the right column lists proteins used for comparisons between X-ray and NMR holo structures. Underlined proteins are shared among the apo and holo categories. Eight proteins showed in *italic* contain multiple holo X-ray structures and multiple NMR conformers. | |

| **Table S4: Multiple X-ray structures and multiple NMR conformers of eight selected proteins.** | | | | | |
| --- | --- | --- | --- | --- | --- |
|  | Protein name  (*Species*) | Uniprot ID | PDB code  X-ray (ligand ID) | Resolution (Å) | PDB code  NMR_conformer ID (ligand ID) |
| 1 | Macrophage metalloelastase  (*Homo sapiens*) | P39900 | 3f17 (hs4)  4gql (r47)  5d2b (56o) | 1.2  1.2  1.2 | 1ycm (ngh)  1ycm_8  1ycm_16 |
| 2 | Fibroblast growth factor 1 (FGF1)  (*Homo sapiens*) | P05230 | 1hkn (n2m)  3k1x (dbx)  3jut (gtq) | 2.0  2.0  2.3 | 1rml (nts)  1rml_18  1rml_3 |
| 3 | Peptidyl-prolyl cis-trans isomerase FKBP1A  (*Homo sapiens*) | P62942 | 1j4h (sub)  1fkf (fk5)  1fkb (rap) | 1.7  1.7  1.9 | 1f40 (gpi)  1f40_2  1f40_8 |
| 4 | UDP-3-O-acyl-N-acetylglucosamine deacetylase  (*Aquifex aeolicus*) | O67648 | 3p3c (3p3)  5u86 (81v)  5drp (5ep) | 1.3  1.6  1.9 | 2jt2 (c90)  2jt2_7  2jt2_22 |
| 5 | E3 ubiquitin-protein ligase MDM2  (*Homo sapiens*) | Q00987 | 4ogn (2u5)  4wt2 (3ud)  4occ (2tz) | 1.4  1.4  1.8 | 2lzg (13q)  2lzg_2  2lzg_3 |
| 6 | Dihydrofolate reductase (DHFR)  (*Homo sapiens*) | P00374 | 1kmv (lii)  3fs6 (dh1)  3ghw ghw) | 1.1  1.2  1.2 | 1yho (trr)  1yho_2  1yho_20 |
| 7 | Collagenase 3  (*Homo sapiens*) | P45452 | 4l19 (1ua)  1xuc (pb3)  2ow9 (sp6) | 1.7  1.7  1.7 | 1fm1 (way)  1fm1_18  1fm1_26 |
| 8 | CREB-binding protein  (CREBP)  (*Homo sapiens)* | Q92793 | 5j0d (69f)  5i86 (69a)  5ktx (6xh) | 1.1  1.1  1.3 | 2l84 (j28)  2l84_11  2l84_18 |
| The table lists eight proteins used for multiple X-ray holo vs multiple NMR holo comparison. Their Uniprot IDs and the corresponding PDB codes, along with the resolution of the X-ray holo structures and NMR conformer IDs from single PDB entry are provided. Ligands determined within holo structures are presented in the brackets. | | | | | |

| **Table S5: Chemical library of database ligands used in docking screens.** | | | | | | | | | | | |
| --- | --- | --- | --- | --- | --- | --- | --- | --- | --- | --- | --- |
|  | Protein name | Database ligand ID | | | | | | | | | |
| Only Apo | Cellular tumor antigen p53 | DB08363 | X0W | UL7 | FY8 | X0U | X0V | QC5 | P74 | KMN | O80 |
|  | Guanyl-specific ribonuclease T1 | GPG | BDBM55013 | BDBM55014 | BDBM52921 | BDBM51785 | BDBM55012 | BDBM51791 | BDBM55010 | BDBM44482 | BDBM50648 |
|  | Replication protein A (70 kDa DNA-binding subunit, REPA) | BDBM260646 | BDBM61188 | BDBM50437278 | BDBM50437283 | BDBM50437285 | BDBM50437299 | BDBM50080584 | BDBM50080693 | BDBM50080769 | BDBM50080820 |
|  | Growth factor receptor-bound protein 2 (GRB2) | BDBM50064333 | BDBM50078348 | BDBM50072862 | BDBM50118687 | BDBM50139762 | DTF | BDBM50072187 | BDBM50241558 | BDBM50080831 | BDBM50161189 |
|  | Heat shock protein (HSP 90-alpha) | 7PP | 94M | 2QA | FU3 | 2GJ | 2R6 | 40W | 40X | 2Q9 | FU5 |
|  | Integrin alpha-L | BQM | BJZ | 803 | AB8 | BDBM50144025 | BDBM50144037 | BDBM50154481 | BDBM50333919 | BDBM50107162 | BDBM50099140 |
|  | Fatty acid-binding protein (FABP) | BDBM50448440 | BDBM50248196 | BDBM50152880 | BDBM50152853 | 75D | BDBM50212876 | BDBM50319700 | BDBM50310988 | 5M8 | BDBM50212876 |
| Holo and Apo | Bcl-2-like protein 1 | BDBM209163 | BDBM209123 | BDBM178583 | BDBM178541 | BDBM178578 | BDBM209147 | BDBM209101 | BDBM209091 | BDBM178690 | BDBM50428721 |
|  | Stromelysin-1 | BDBM50099857 | NGH | OHL | ATT | 0DS | HQQ | S27 | BDBM50063917 | BDBM50076995 | BDBM50031798 |
|  | Macrophage metalloelastase | BDBM50180609 | 068 | 077 | PF3 | EEA | KLG | BDBM50216005 | BDBM50295499 | 37A | BDBM50270675 |
|  | Fibroblast growth factor 1 (FGF1) | BDBM50143013 | BDBM50144520 | BDBM50144523 | BDBM50144525 | BDBM50097125 | BDBM50143019 | BDBM50143025 | BDBM50143012 | BDBM50422562 | BDBM50143009 |
|  | Peptidyl-prolyl cis-trans isomerase FKBP1A | BDBM50228108 | BDBM50228117 | BDBM50228111 | BDBM23347 | BDBM50068572 | BDBM50113058 | BDBM500695576 | BDBM50068573 | BDBM50068608 | BDBM50086090 |
| Only Holo | UDP-3-O-acyl-N-acetylglucosamine deacetylase | BDBM50115452 | BDBM92271 | BDBM92260 | BDBM92272 | BDBM92283 | BDBM92280 | BDBM92263 | BDBM50115463 | BDBM92259 | BDBM92255 |
|  | E3-ubiquitin ligase MDM2 | 4T4 | MI6 | NUT | Y30 | 0R3 | 28W | 4NJ | 7HC | 2SW | 35T |
|  | Dihydrofolate reductase | 684 | COP | TOP | OAG | MXA | MTX | 1QZ | 1XF | 9DR | 3TU |
|  | Collagenase 3 | 302 | MSB | 24F | BDBM50142473 | 3EK | 3EJ | 3KE | RS1 | BDBM50160843 | HS5 |
|  | CRE-binding protein (CREBP) | UL4 | 6XG | 6XB | 2O4 | TTR | L85 | 3PF | 2LO | 68Y | 98 |
|  | E3 ubiquitin ligase XIAP | SMK | BDBM26218 | X23 | CO9 | G13 | 419 | BDBM44323 | BDBM50436874 | BDBM50302896 | BDBM5039363 |
| Protein name, name of the proteins used in this study; Database ligand ID, IDs of 10 experimentally validated ligands. Ligands derived from the BindingDB consist of BDB code followed by a number, while ligands derived from the PDB are consisted of three letter codes. | | | | | | | | | | | |

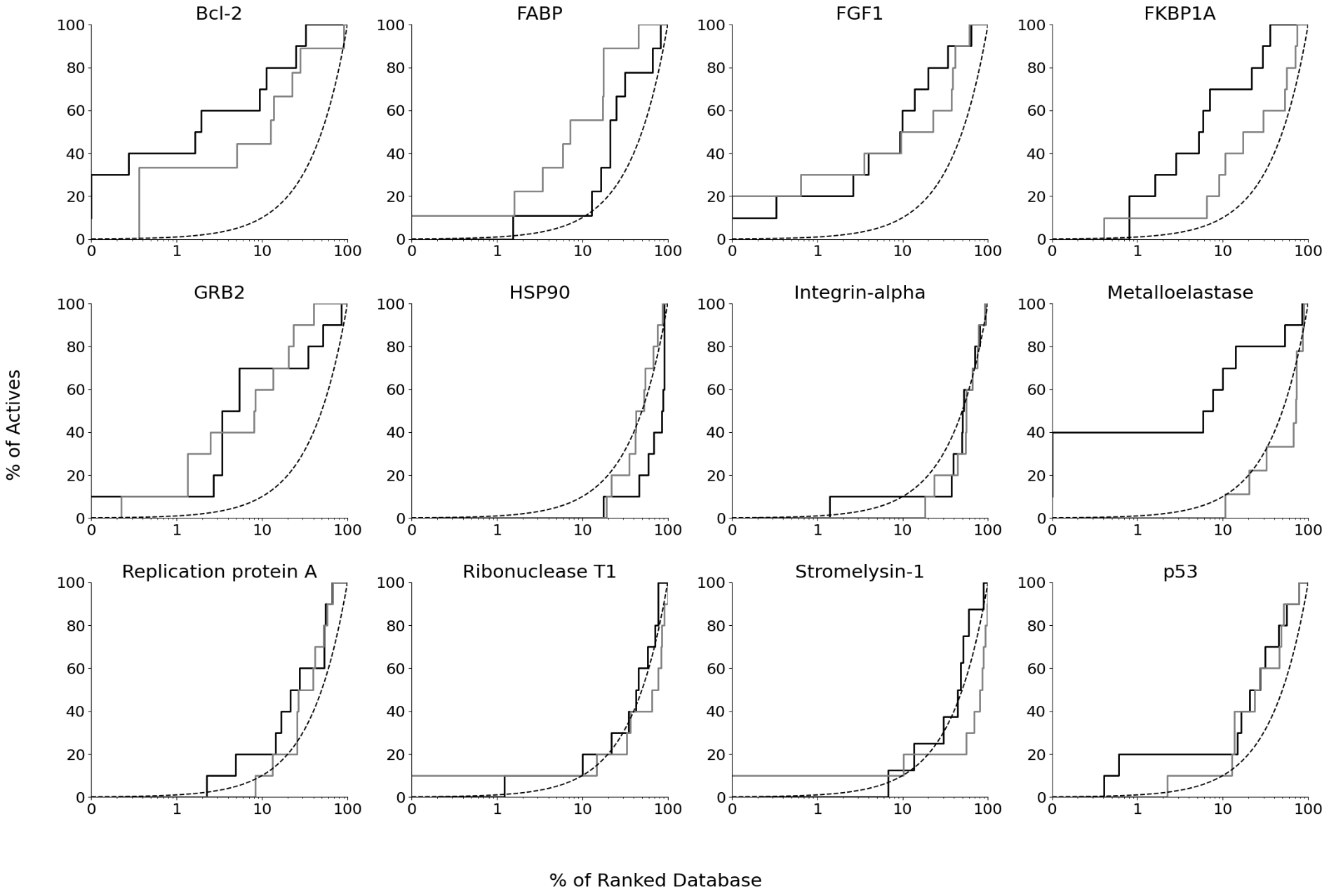

**Figure S1. Enrichment curves of apo X-ray/NMR structures.** Enrichment plots are plotted for X-ray apo structures (black), NMR apo structures (grey), and random selection (dotted line).

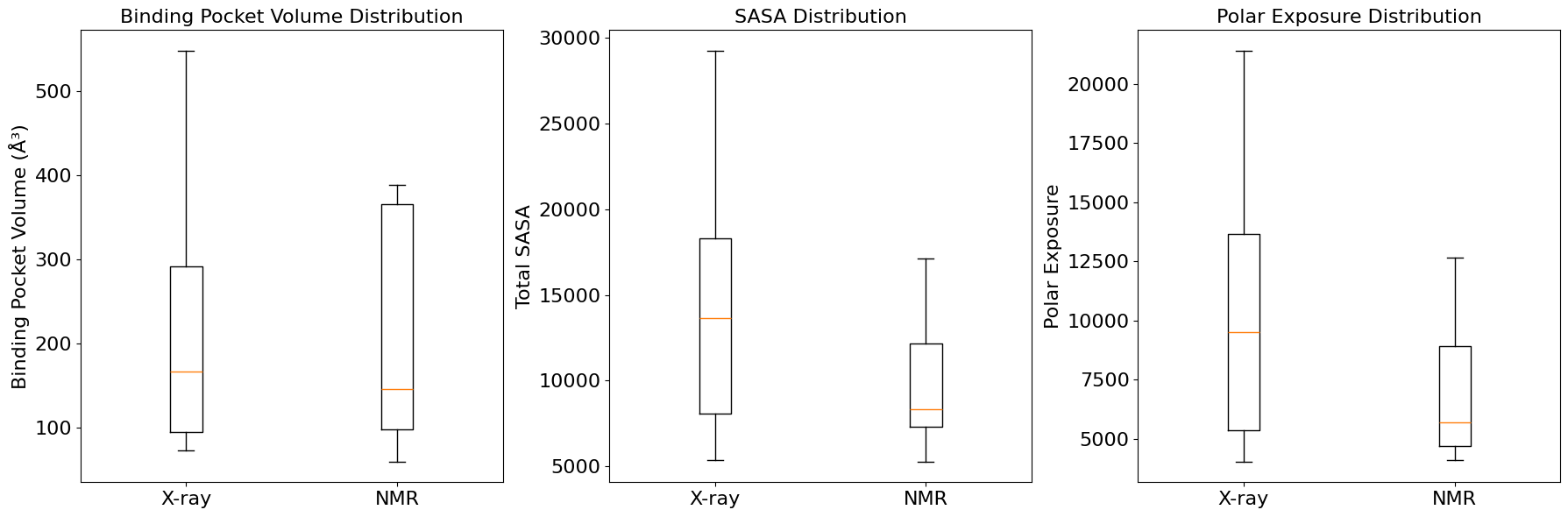

**Figure S2. Box plots comparing structural properties of apo proteins solved with X-ray crystallography and NMR spectroscopy.** Binding pocket distribution volume distribution (Å^3^) shows the deviation in pocket size between X-ray and NMR structures (Left). SASA distribution indicates differences in overall solvent exposure to the solvent (Middle), while Polar exposure distribution represents the extent of polar regions exposed to the solvent. These distributions highlight systematic variations between structural determination methods, with X-ray structures generally exhibiting larger values across all three metrics.

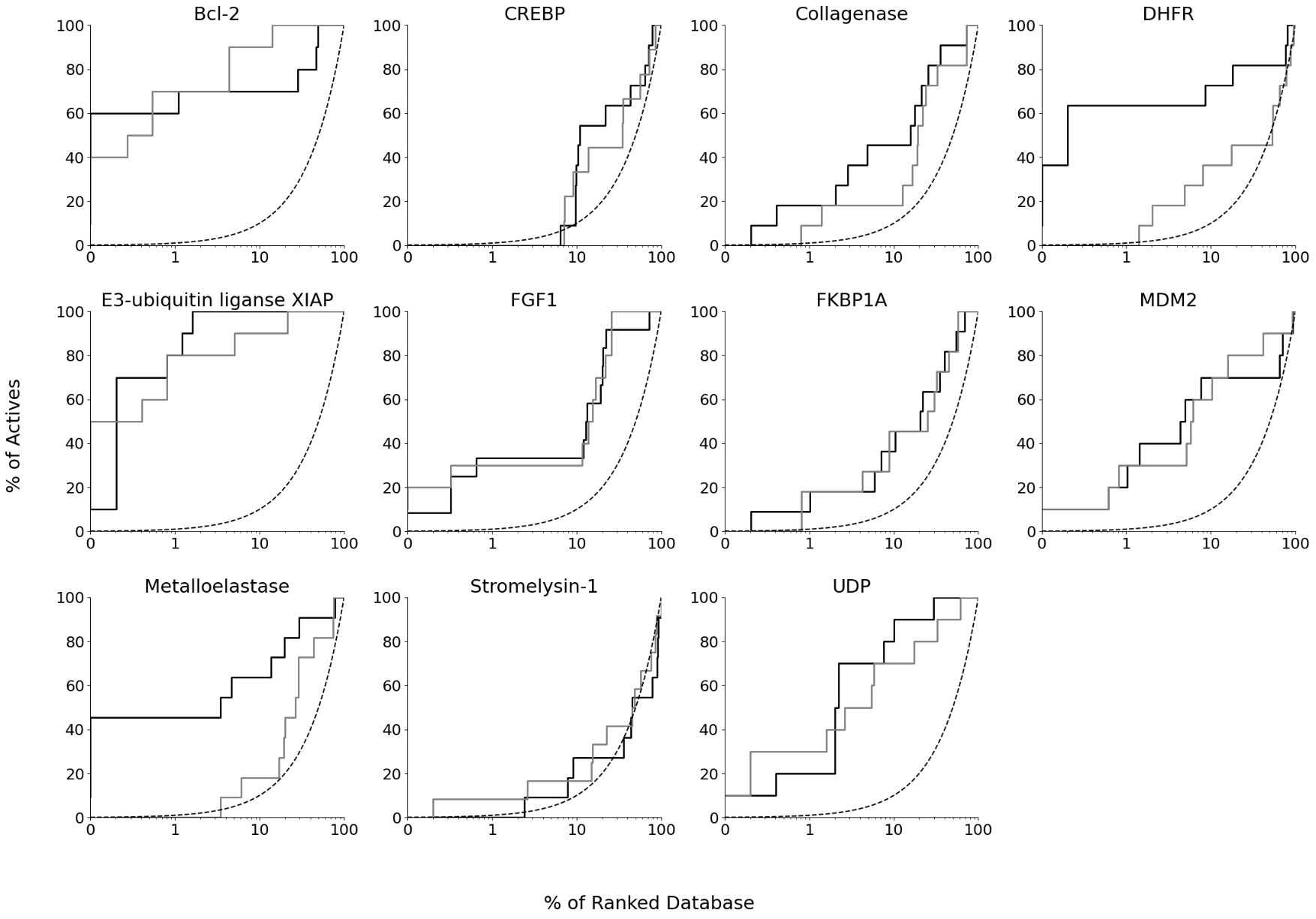

**Figure S3. Enrichment curves of single holo X-ray/NMR structures.** Enrichment plots are plotted for X-ray holo structures (black), NMR holo structures (grey), and random selection (dotted line).

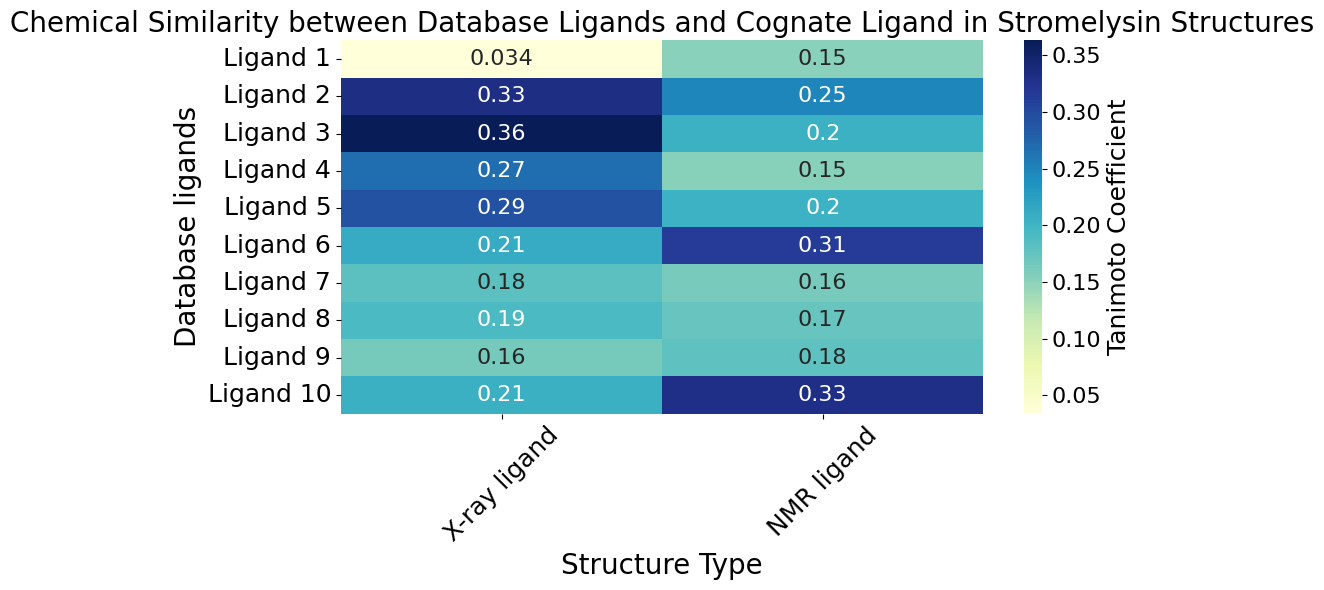

**Figure S4. Comparison of chemical similarity between Stromelysin X-ray/NMR cognate ligand and database ligands used in molecular docking.**

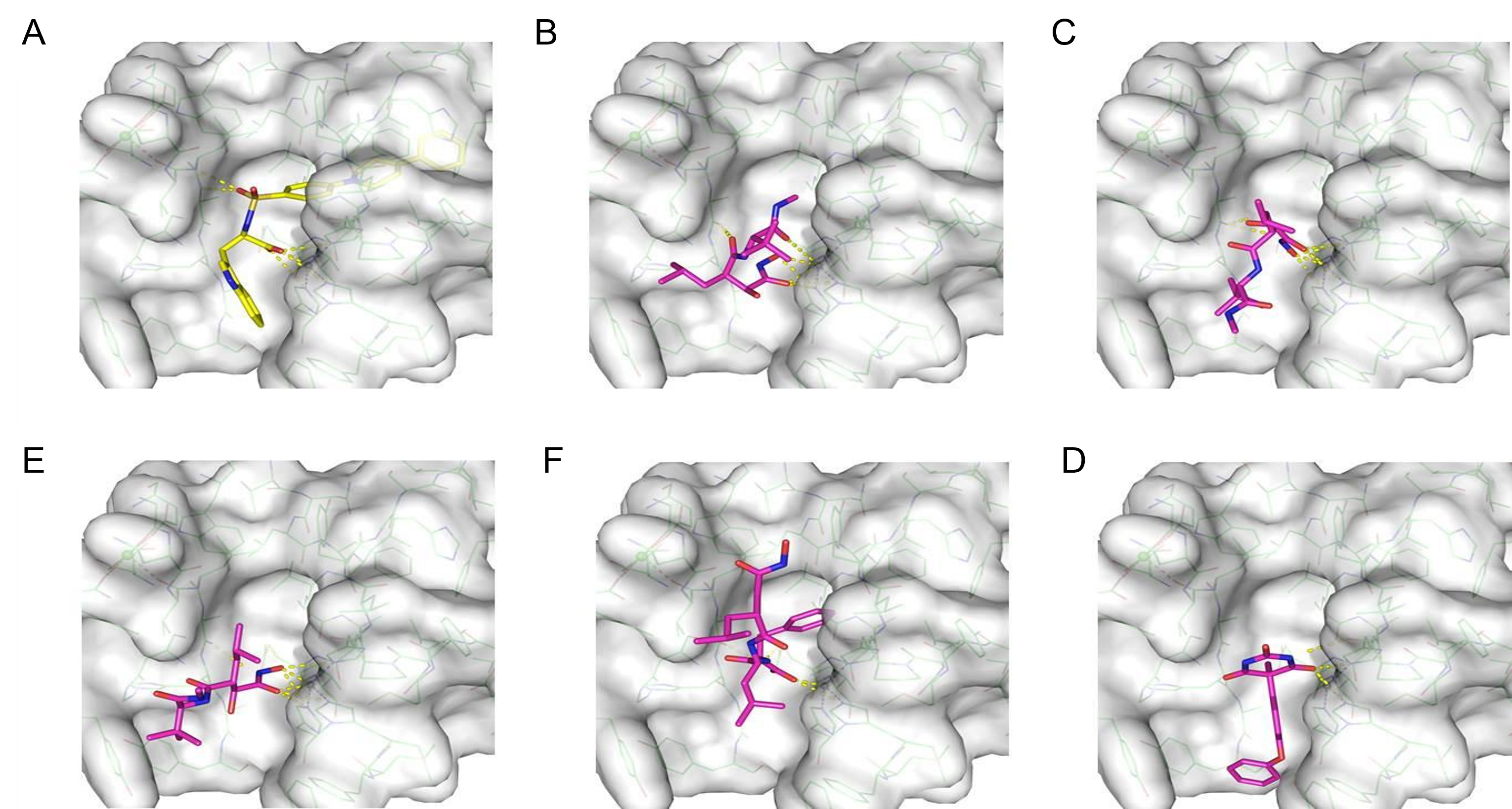

**Figure S5**. **Binding poses of top-ranked ligands used in virtual screening over the X-ray holo structure of Stromelysin-1 (PDB ID: 1CIZ)**. The linear conformation of the crystal ligand (PDB ligand ID: DPS) (A, yellow stick) fits snugly within the tunnel-like binding pocket, achieving a high docking score of -12.22. In contrast, the docked ligands—B: OHL (-8.45), C: 0DS (-7.32), D: ATT (-6.90), E: S27 (-6.47), and F: HQQ (-5.95) (magenta stick)—remain partially solvent-exposed, resulting in lower docking scores. This underscores the better accommodation of the cognate ligand (A) compared to the virtual screening ligands (B-F), which fail to fully occupy the binding pocket.

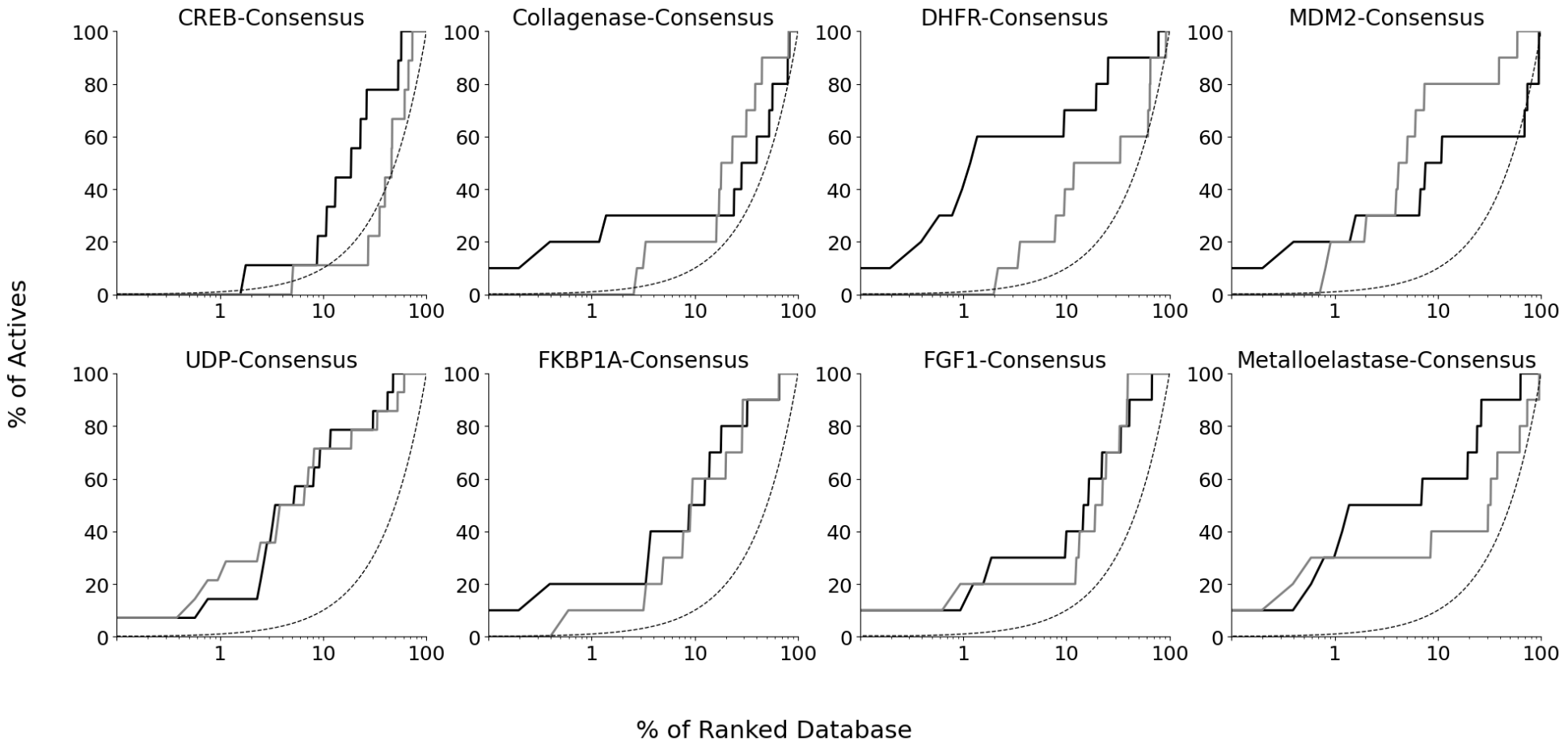

**Figure S6. Enrichment curves of consensus holo X-ray/NMR structures.** Enrichment plots are plotted for X-ray holo structures (black), NMR holo structures (grey), and random selection (dotted line).

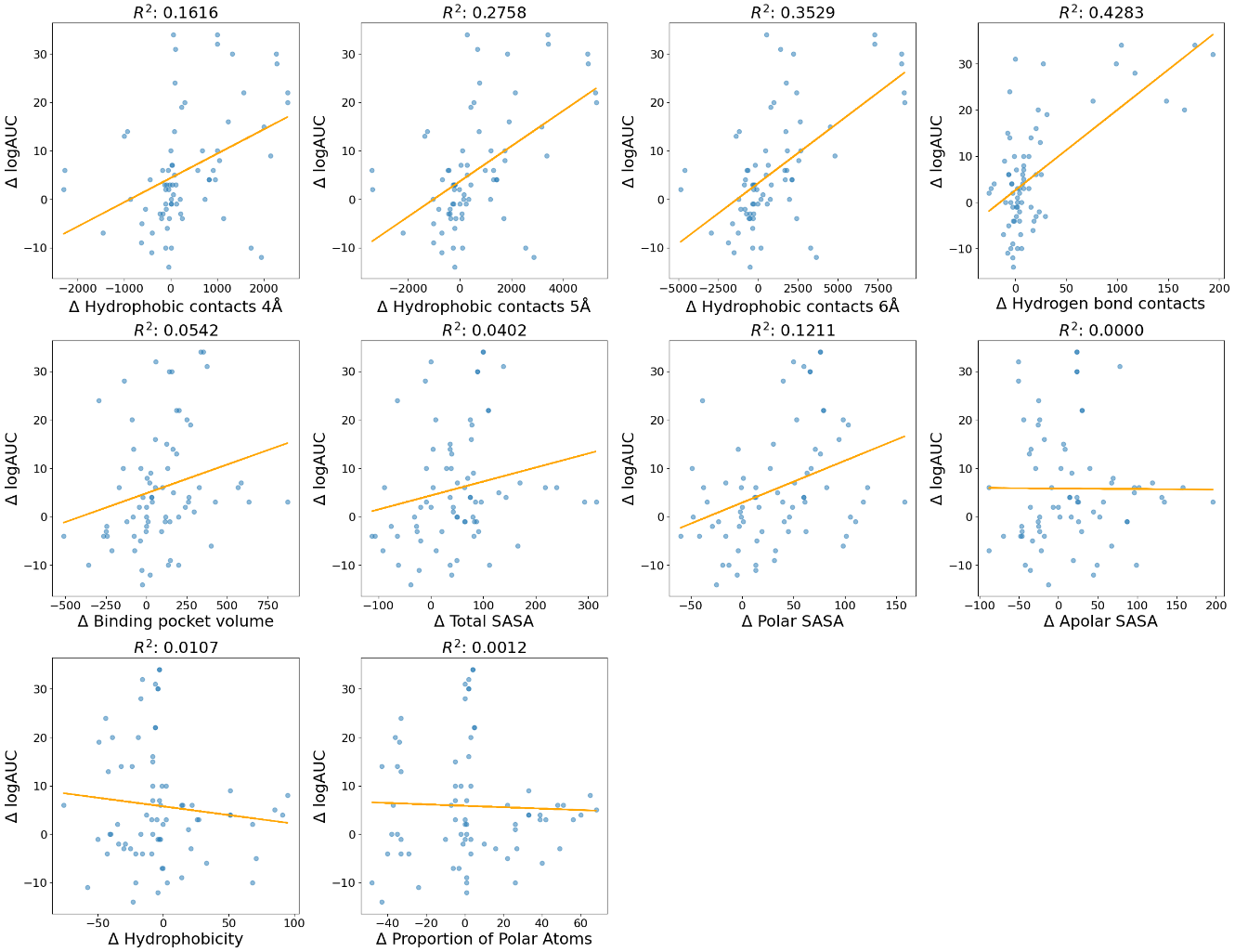

**Figure S7. Scatter plots illustrating the correlation between differences in binding site and protein-ligand interaction features, and virtual screening performance (Δ logAUC).** The top row shows correlations between Δ logAUC and the number of hydrophobic contacts formed at various distances (4Å, 5Å, and 6Å), as well as the number of hydrogen bonds, with R² values of 0.16, 0.28, 0.35, and 0.43, respectively, indicating moderate correlations. The middle and bottom rows depict correlations between Δ logAUC and binding pocket volume, total SASA, polar SASA, hydrophobicity, and polar atom proportion, all of which show low or negligible correlation (R² values ranging from 0.00 to 0.13).

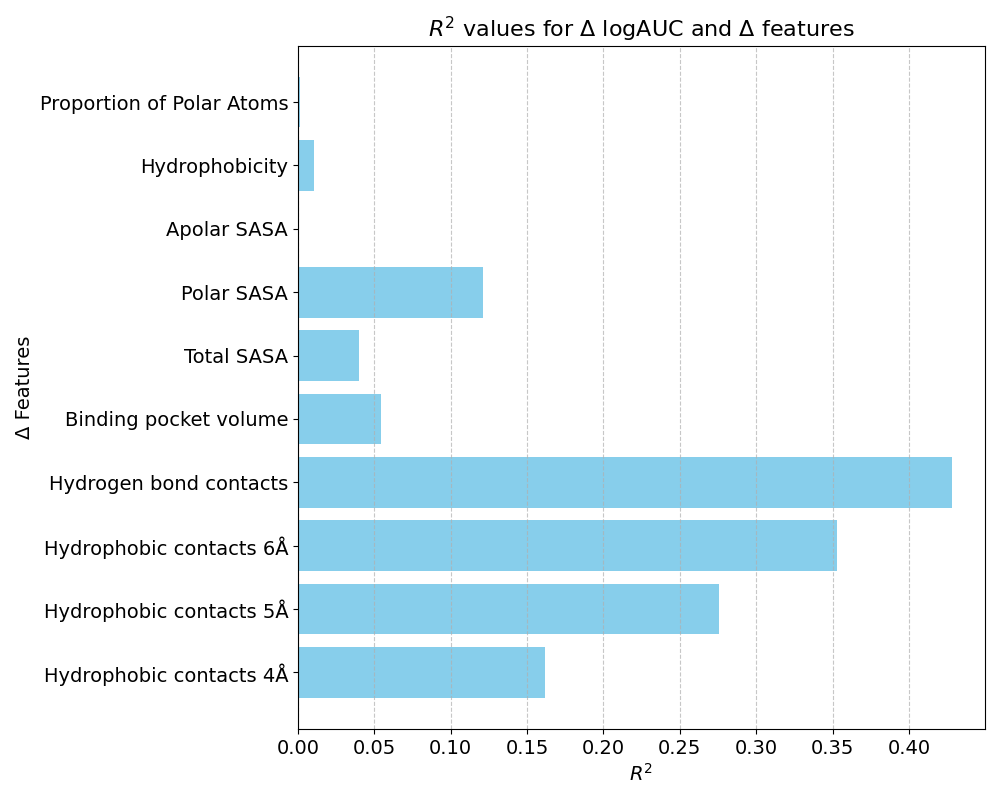

**Figure S8. Bar chart showing the R² values for the correlation between differences in protein-ligand interaction and binding site features, and virtual screening performance (Δ logAUC)**. The features with the highest correlation are hydrophobic contacts at 4Å, 5Å, and 6Å distances, with R² values of 0.16, 0.28, and 0.35, respectively. Hydrogen bonds also exhibit a relatively strong correlation (R² = 0.43). In contrast, binding pocket volume, total SASA, and polar SASA show weaker correlations (R² values between 0.0 and 0.13), while hydrophobicity, apolar SASA, and polar atom proportion exhibit negligible correlation.

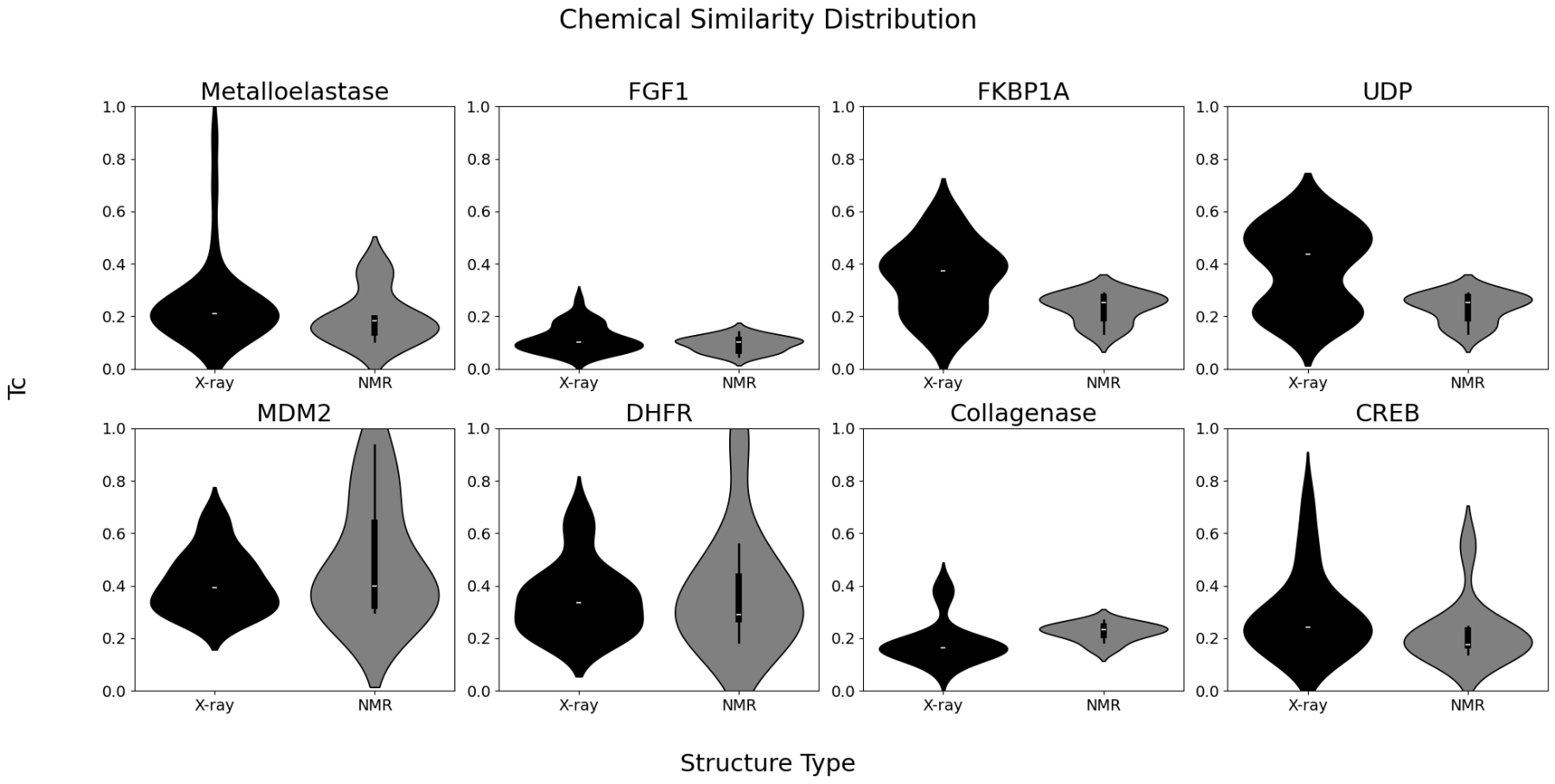

**Figure S9. Violin plots depicting the distribution of chemical similarity (Tc) values calculated between cognate ligands of X-ray/NMR structures and database ligands used in molecular docking.** Each subplot corresponds to a single protein, comparing Tc value distributions across X-ray (black) and NMR (grey) structures. The width of each violin represents the KDE density of values, with white markers indicating median Tc values. Across most proteins, cognate ligands of X-ray and NMR holo structures show distinct chemical similarities to the ligands used in molecular docking.
